# A single mutation in the ‘DSL’ motif of acyl carrier protein can prevent its *in vivo* modification by *E. coli* Holo-acyl carrier protein synthase (AcpS)

**DOI:** 10.1101/2025.01.28.635281

**Authors:** Chetna Dhembla, Debodyuti Sadhukhan, Rashima Prem, Shivangi Vaish, Shalini Verma, Suman Kundu, Monica Sundd

## Abstract

*E. coli* expression system is the method of choice to obtain high yields of a pure protein. However, there is always a chance that an overexpressed protein shares structural or sequence homology with the substrate of an *E. coli* enzyme. In such cases, the expressed protein may be partially or fully converted into the product. A notable example is the expression of acyl carrier proteins (ACP) in *E. coli*. Since most type II ACPs of the fatty acid synthesis pathway (FAS) have a conserved helix II, the carrier proteins are recognized as a substrate by Holo-acyl carrier protein synthase (AcpS). Thus, most ACPs express as partially or fully loaded proteins in *E. coli*. This undesirable modification is a concern when the objective is to obtain milligram amounts of apo-ACP. Here, using an approach combining mutagenesis, enzyme activity, and NMR, we probed for residues in ACP that can prevent this *in vivo* modification, without affecting Sfp (Surfactin synthetase activating enzyme) function. Taking cues from the *E. coli* ACP-AcpS structure (PDB 5VCB), charge neutralization mutations were designed at five different positions in *Ec*ACP that participate in ion-pair interaction with AcpS. Three of the mutants expressed solely as apo-ACP in *E. coli viz.* D35N, E41A and E47A/E48A. However, only the D35N mutant could be converted into holo-/acyl-ACP using Sfp *in vitro*, establishing mutagenesis as a viable strategy to prevent undesired modifications *in vivo*. As proof of principle, the mutation was applied to two unrelated ACPs that express primarily as modified proteins in *E. coli* -*Mus musculus* mitochondrial FAS ACP (*Mm*ACP) and *Salmonella* Typhimurium invasion acyl carrier protein (IacP). Single D35N mutation of the ACPs prevented their *in vivo* modification by AcpS, and the mutants were efficiently converted into holo-ACP by Sfp *in vitro*. These results demonstrate that D35N mutagenesis is a useful strategy to express apo-ACP in *E. coli* and is applicable across all type II ACPs. Furthermore, we show that holo-IacP and holo-*Mm*ACP are not recognized as substrates by AcpH (*E. coli* Acyl carrier protein hydrolase), and therefore they express predominantly as modified proteins in *E. coli*.

## INTRODUCTION

Acyl carrier proteins (ACP) play an essential role in the biosynthesis of fatty acids, polyketides, non-ribosomal peptides, and other bioactive molecules (*1, 2*). They exist as free proteins, or as components of a large, multidomain, multifunctional complex. ACPs are expressed as apo-proteins (inactive form) and are post translationally modified into their active holo-form by the catalytic action of a 4’-phosphopantetheinyl transferase(*3*). In this process, a conserved serine of the ACP molecule is posttranslationally modified by the attachment of a coenzyme A derived 4’-phosphopantetheinyl moiety, resulting in the formation of holo-ACP. Three groups of phosphopantetheinyl transferases (PPTases)-Group I, II and III catalyze this reaction, classified based on structural organization (*4, 5*). Most group I enzymes are typically homotrimers, comprising three identical subunits of approximately 110 amino acids, with an α+β fold. The functional enzyme has three active sites, each located in the cleft between the two monomers. These enzymes display a broad substrate specificity and are known as holo-acyl carrier protein synthase (AcpS). Group I phosphopantetheinyl transferases modify ACPs involved in primary metabolism, *i.e.* fatty acid synthesis pathway of *E. coli, Bacillus subtilis, M. tuberculosis, Streptococcus pneumoniae, etc.* (*6, 7*). Group II PPTases are monomeric, consisting of an N-and a C-terminal α+β fold domains, and are approximately 220 amino acids in length. Each domain is structurally equivalent to a subunit of the Group I PPTase. The active site is located at the junction of the two domains, and the molecule has a pseudo-2-fold symmetry. PPTases belonging to group II have a broader substrate specificity than Group I, and activate ACPs involved in secondary metabolism, such as nonribosomal peptide synthesis. A well-known example from this group is Surfactin synthetase activating enzyme (Sfp) from *B. subtilis*, which post-translationally phosphopantetheinylates seven peptidyl carrier protein domains of surfactin synthetase (*8, 9*). Group II PPTases are found in *Bacillus subtilis, Homo sapiens, Streptomyces tsukubaensis etc.* (*10, 11*). Group III PPTases exist as domains of a large, multidomain, multifunctional fatty acid/polyketide synthesis machinery. They phosphopantetheinylate ACP domains prior to the assembly of the megacomplex and are highly specific for their cognate ACP domains. Group III PPTases are found in *Saccharomyces cerevisiae*, *Candida albicans, Mycobacterium tuberculosis, etc.* (*12*).

Interestingly, the phosphodiester bond between the -OH group of the conserved Ser of an ACP and the phosphate group (PO4^-^) of a phosphopantetheine can be cleaved by an acyl carrier protein phosphodiesterase, also known as acyl carrier protein hydrolase (AcpH, PDB ID 1TIK). AcpH functions in a direction opposite to PPTase, converting holo-or acylated-ACPs into apo-ACP. An *E. coli* strain harboring the AcpH deletion mutant grew normally, suggesting that the enzyme is not essential for survival. However, AcpH is necessary for recycling/regenerating the prosthetic group, for efficient bacterial growth (*13*). Akin to AcpS, acyl-ACP hydrolase (AcpH) also has a broad substrate specificity and can hydrolyze serine linked phosphopantetheine moiety (4’-PP) either alone or when tethered to a C_6_-C_16_ carbon chain. Limited information exists on the precise substrate specificity of AcpH (*13*).

Since most type II ACPs share a very similar four helix bundle fold, a conserved helix II, and a ‘DSL’ motif, Group I & Group II PPTases often phosphopantetheinylate non-cognate ACPs as well. This promiscuity becomes a major drawback when a recombinant ACP is expressed in *E. coli*. Due to the presence of a group I PPTase (AcpS) endogenously, that has a broad substrate specificit*y*, a substantial amount of the ACP is expressed in the phosphopantetheinylated form. Notably, the extent of modification varies from one ACP to the other. For instance, some type II ACPs, such as *Leishmania* ACP are expressed as unmodified proteins, while *E. coli, P. falciparum,* and *M. tuberculosis* ACP display partial modification. In a few other cases, such as human mitochondrial ACP (*Hs*ACP), the protein is expressed primarily in the phosphopantetheinylated form (*14*). Since the phosphopantetheinylated ACPs are also recognized as substrates by *E. coli* fatty acid synthesis machinery, multiple acylated forms may also be present (*15*). As a result, additional purification steps are required to separate the apo-protein from the holo-and acylated forms, resulting in lower yields of the apo-protein. With the intent to prevent this unwanted modification of non-native ACPs in *E. coli*, the present study was initiated. Our studies demonstrate that a charge neutralization mutation (D35N) in the ‘DSL’ motif of ACP can completely block this AcpS catalyzed modification *in vivo*, allowing higher yields of the apo-protein. Importantly, this single mutation preserves the structural and functional features of ACP and does not interfere with Sfp function. Altogether, we show that the extent of ACP modification is determined by the collective action of two *E. coli* enzymes, a) holo-acyl carrier protein synthase (AcpS), and b) acyl carrier protein hydrolase (AcpH), that work in opposite directions.

## MATERIALS AND METHODS

### Cloning, expression and purification

*Salmonella* Typhimurium genome was obtained from BEI resources, USA. The gene encoding *Mus musculus* mitochondrial ACP was synthesized after codon optimization. PCR amplification of S. Typhimurium (LT2 strain) invasion acyl carrier protein (UniProtKB P0CL00), and *M. musculus* mitochondrial ACP (UniProtKB Q569N0) genes was performed by introducing NdeI and HindIII restriction sites in the forward and reverse primers, respectively. The resulting PCR products were cloned in a pET28a(+) vector and the inserts were verified by sequencing.

*E. coli* Acyl carrier protein phosphodiesterase (UniProtKB P21515) was cloned from the *E. coli* genome and expressed as an MBP fusion protein with an N-terminal 6X his tag. *Leishmania major* ACP (*Lm*ACP, UniProtKB E9AD06) and *E. coli* AcpP (UniProtKB P0A6A8) were previously cloned in the lab (*16*).

For protein expression, plasmids carrying the desired gene were transformed into *E. coli* BL21(DE3)/Rosetta (DE3) cells. The transformed cells were plated on LB Agar plates supplemented with kanamycin sulfate (final concentration 50 µg/ml). After overnight incubation at 37 °C, a single colony was inoculated in a 10 ml LB broth (supplemented with kanamycin sulfate). The culture was kept at 37 °C with constant shaking at 180 rpm. The starter culture was then inoculated into a 1 litre autoclaved LB broth (supplemented with 50 µg/ml kanamycin sulfate) and incubated at 37 °C, with constant shaking at 180 rpm, till the optical density (A_600_) reached 0.8-1.0. Protein overexpression was induced by the addition of 0.25-0.5 mM IPTG, followed by incubation at 25 °C (overnight), with constant shaking at 180 rpm. Bacterial cells were harvested by centrifugation at 5000g for 30 minutes at 4°C, and stored at -20 °C.

To purify His-tagged proteins, the bacterial cell pellet was resuspended in 50 mM Tris-HCl (pH 7.8), 150 mM NaCl (lysis buffer) and lysed by sonication. ACP phosphodiesterase treatment was performed using the endogenous enzyme by adding 10 mM MgCl_2_ and 2 mM MnCl_2_ to the lysate, and incubating at 35°C for 2 hour at 120 rpm to convert holo-ACP into apo-ACP. The lysate was centrifuged at 15,000g for 30 min at 4°C. The supernatant was applied to a Ni^2+^-NTA column pre-equilibrated with the resuspension buffer (50 mM Tris-HCl buffer (pH 7.8), 150 mM NaCl) at room temperature. The column was washed with 100 ml resuspension buffer containing 10 mM imidazole to remove nonspecifically bound proteins. The target protein was eluted using an imidazole gradient (10-250 mM) in the same buffer. Eluted fractions were analyzed by 12% SDS-PAGE gel, and the pure fractions were pooled and concentrated using a 3 kDa (15ml) Amicon Centricon. The his-tag present at the N-terminus of the protein was cleaved using immobilized thrombin. For *Mm*ACP, an additional step of 50% isopropanol treatment was used to precipitate the nonspecificially bound *E. coli* Eftu (Elongation factor temperature unstable 2) that copurifies with mitochondrial ACPs (*14*).

In cases where the ACP was expressed in both apo-as well as holo-forms, an additional step of anion-exchange chromatography was introduced using AKTA resource Q column. Prior to loading the sample onto the column, the protein was buffer exchanged into 20 mM sodium phosphate (pH 6.0), and the same buffer was used for ion exchange chromatography. The two forms of ACP eluted separately based on charge under a 0-500 mM NaCl gradient. The pure fractions were pooled, concentrated, and loaded onto an Akta Hiload 16/60 Superdex 75 column. The pure fractions were once again pooled, concentrated, and stored at -20 °C. The purity of the protein was assessed using 12% Native-PAGE, 12% SDS-PAGE, C_18_-reverse phase chromatography, and in some cases, ESI Mass Spectrometry (*17*).

*Bacillus subtilis* Sfp (UniProtKB P39135) was expressed in *E. coli* Bl21(DE3) cells using a previously described protocol (*16*).

### Site directed mutagenesis

Single or double mutations were introduced in ACP using site-directed mutagenesis approach, and the resulting mutants were purified analogous to the wild type protein.

### Electrospray Ionisation Mass Spectrometry (ESI-MS) analysis

Wild-type ACPs and their mutants were concentrated and desalted using C_4_ or C_8_ ZipTips. Protein bound to the ZipTip was eluted in 50 % acetonitrile, 0.1 % formic acid, and subjected to ESI-MS analysis. Data were visualized using Thermo Scientific Xcalibur software.

### NMR sample preparation and data acquisition

Uniformly labeled ^1^H^15^N^13^C wild-type *Ec*ACP and mutants were overexpressed by growing *E. coli* BL21(DE3) cells in M9 minimal media containing ^15^N ammonium chloride as the nitrogen source. The media was autoclaved, and after cooling, supplemented with sterile filtered 0.4% ^13^C or ^12^Cglucose, 2 mM MgCl_2_, 20 µM CaCl_2_, vitamins and appropriate antibiotics. Protocol for induction, overexpression, and purification was the same as the unlabeled protein.

For NMR titrations, the ACP sample comprised 0.15-0.3 mM protein in 50 mM Tris-HCl buffer (pH 7.8), 150 mM NaCl, and 10 % D_2_O. For ^1^H^15^N HSQC experiments with the mutants, 0.3-0.5 mM ACP sample was prepared in 20 mM Sodium Phosphate buffer (pH 7.0), 100 mM NaCl. All NMR data were recorded at 293 K on a Bruker Avance III 700 MHz NMR spectrometer, equipped with a TXI/QCI probe, at the National Institute of Immunology (New Delhi, India). NMR data were processed on a workstation running Red Hat Enterprize Linux 5.0, using NMRDraw/NMRPipe (*18*), and analyzed with Sparky (*19*). The data were multiplied by a phase shifted sine bell apodization function in all dimensions.

^1^H^15^N HSQC spectra were acquired with 2048 data points in the t2 dimension and 128-512 data points in the t1 dimension. The data were linear predicted in the forward direction for up to half the number of experimental points in the indirect dimensions. ^15^N^13^C spectra were referenced indirectly using sodium 2, 2-dimethyl-2-silapentane-5-sulfonate (DSS) as a chemical shift standard (*20*).

### Conversion of apo-to holo-/acyl-ACP using Sfp

Apo-ACP was converted into holo-and acyl-ACP by incubating 20 μM apo-ACP, 5 μM *Bacillus subtilis* 4’-Phosphopantetheinyl transferase (Sfp), and 100 μM acyl-/CoA in 50mM Tris-HCl (pH 8.0), and 2 mM MgCl_2_, using a modified protocol published previously (*21*). The assay samples were incubated at 37 °C for 2 hours. The conversion was monitored by 12% Native-PAGE gel, as the holo-form migrates ahead of the apo form due to higher negative charge imparted by the PO_4_ ^-^. The results were confirmed by C_18_-HPLC reverse phase chromatography, and ESI-MS. For NMR samples, the reaction volume was scaled up accordingly.

### Acyl carrier protein hydrolase (AcpH) assay to convert holo-to apo-ACP

An *in vitro* reaction mixture containing 20 µM holo-ACP, 50 mM Tris-HCl (pH 8.0), 5 mM MgCl_2_, 200 µM MnCl_2_, 2 mM DTT and 10 µM AcpH enzyme was prepared in a final volume of 20 µl. The reaction products were analysed by 12% Native-PAGE (*13*).

## RESULTS

### Acyl carrier proteins exhibit varying levels of phosphopantetheinylation when expressed in *E. coli*

*E. coli, P. falciparum and M. tuberculosis* type II acyl carrier proteins (ACP) express in two different forms: apo-and holo-ACP when expressed in *E. coli.* In contrast, human mitochondrial ACP is expressed primarily as a phosphopantetheinylated /acylated protein (*14, 15, 21–23*). To understand the molecular basis for this disparity, four type II ACPs were selected: *E. coli* (*Ec*ACP), *Leishmania major* (*Lm*ACP), mitochondrial *Mus musculus* (*Mm*ACP) and Salmonella Typhimurium invasion acyl carrier protein (IacP). The first three proteins (*Ec*ACP, *Lm*ACP, and *Mm*ACP) are canonical type II ACPs involved in the fatty acid synthesis pathway. However, IacP is an atypical/specialized ACP, required for the acylation of the cell invasion protein SipB (*Salmonella* invasion protein B), which participates in the type III secretion system (T3SS) of Salmonella Typhimurium. Both *Lm*ACP and *Mm*ACP are mitochondrial proteins, expressed with an N-terminal mitochondrial targeting sequence. However, the recombinant mitochondrial proteins used in this study were expressed without the signal peptide. Figure 1A shows a sequence alignment of the four types II ACPs used in this study. Identical residues in the four proteins are highlighted with a black background, while the residues identical to *Ec*ACP but not conserved across all four ACPs are shown in grey. Sequence comparison of the four ACPs suggests conservation in the loop I and helix II region.

**Figure 1.**
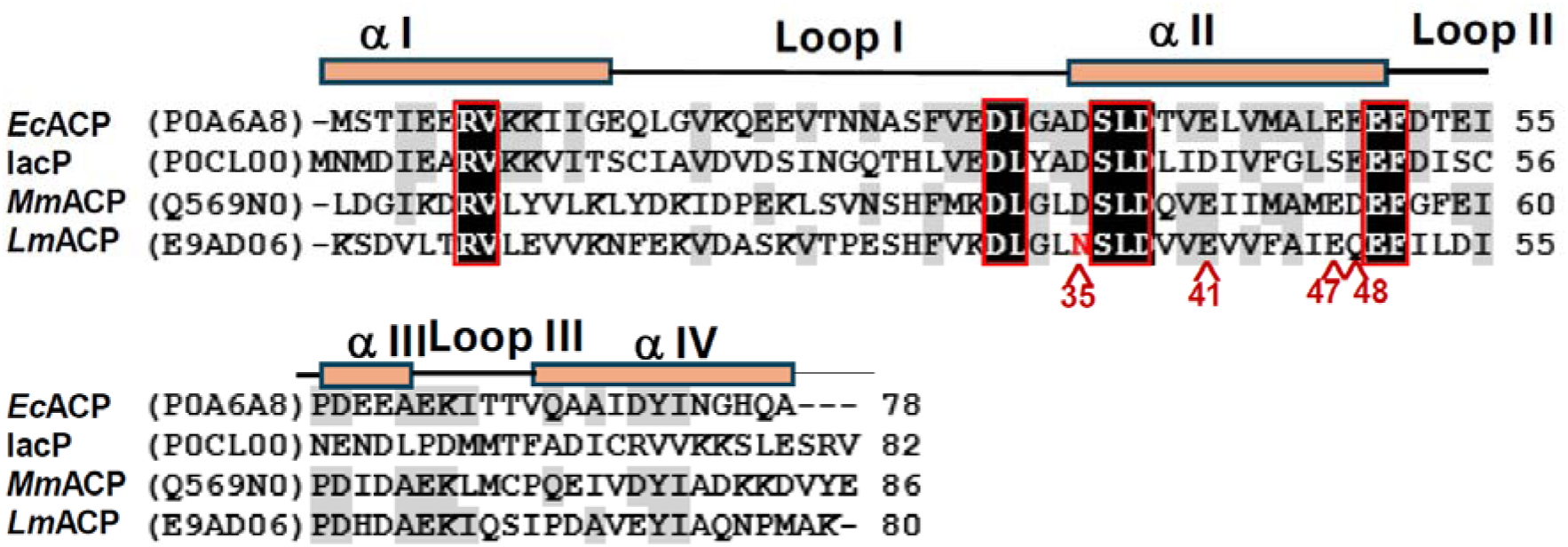
Sequence comparison of type II ACPs from different sources. The amino acid sequence of *E. coli* ACP (*Ec*ACP, UniProtKB P0A6A8), aligned with the Invasion acyl carrier protein (IacP, UniProtKB P0CL00), mitochondrial *Mus musculus* ACP (*Mm*ACP, UniProtKB Q569N0), and *Leishmania major* ACP (*Lm*ACP, UniProtKB E9AD06). Sequence alignment was performed using Clustal Omega (*24*). Residues identical in all four sequences are shown with a black background, while the ones identical with *Ec*ACP but not across all four ACPs are highlighted in grey. Amino acids mutated in this study are marked and listed at the bottom in red.

The four proteins were purified using Ni^2+^-affinity chromatography and analyzed by 12% Native-PAGE. Purified *Lm*ACP, IacP and *Mm*ACP displayed a single band on native-PAGE, indicating a single form of ACP, which could be either apo-or phosphopantetheinylated. In contrast *Ec*ACP expressed in two forms, apo-and holo-, which were subsequently purified using anion-exchange chromatography.

Since *Mm*ACP, *Lm*ACP, and IacP expressed in only one form, it was necessary to understand whether the expressed proteins correspond to the apo-or the holo-form. Therefore, enzymatic conversion of the three ACPs was carried out using Sfp (*Bacillus subtilis* type II PPTase that has a broad substrate specificity) and CoA/C_8_-CoA. Control samples (without Sfp) loaded on a 12% Native-PAGE are shown for *Lm*ACP (lane#3), IacP (lane#5) and *Mm*ACP (lane#7) in Fig. 2A. For comparison, Sfp assay was carried out with apo-*Ec*ACP as well (as a positive control). A notable increase in the migration of the ACP band after the assay was considered as an indicator of holo-ACP formation. The protein bands in the assay samples for *Ec*ACP (lane #2) and wt*Lm*ACP (lane #4) displayed faster migration on native-PAGE compared to their respective control bands shown in lanes #1 & #3, respectively. The results suggest that both the proteins were in the apo-form in the control sample and were converted into the holo-form during the Sfp assay. Relatively less conversion was observed in *Lm*ACP, as the protein required longer than 2 hours to fully convert into holo-ACP (*17*). In contrast, Sfp assay for wild type IacP (lane #6) and *Mm*ACP (lane #8) incubated for the same duration, did not display any change in migration compared to control, shown in lanes # 5 & #7, respectively. The *Mm*ACP assay was performed using both coenzyme A and C_8_-CoA as substrates, but the gel bands are shown only for the samples incubated with C_8_-CoA in Fig. 2A. Based on the pattern of migration, it is apparent that no modification occurred in *Mm*ACP and IacP samples when incubated with Sfp and CoA/C_8_-CoA.

**Figure 2.**
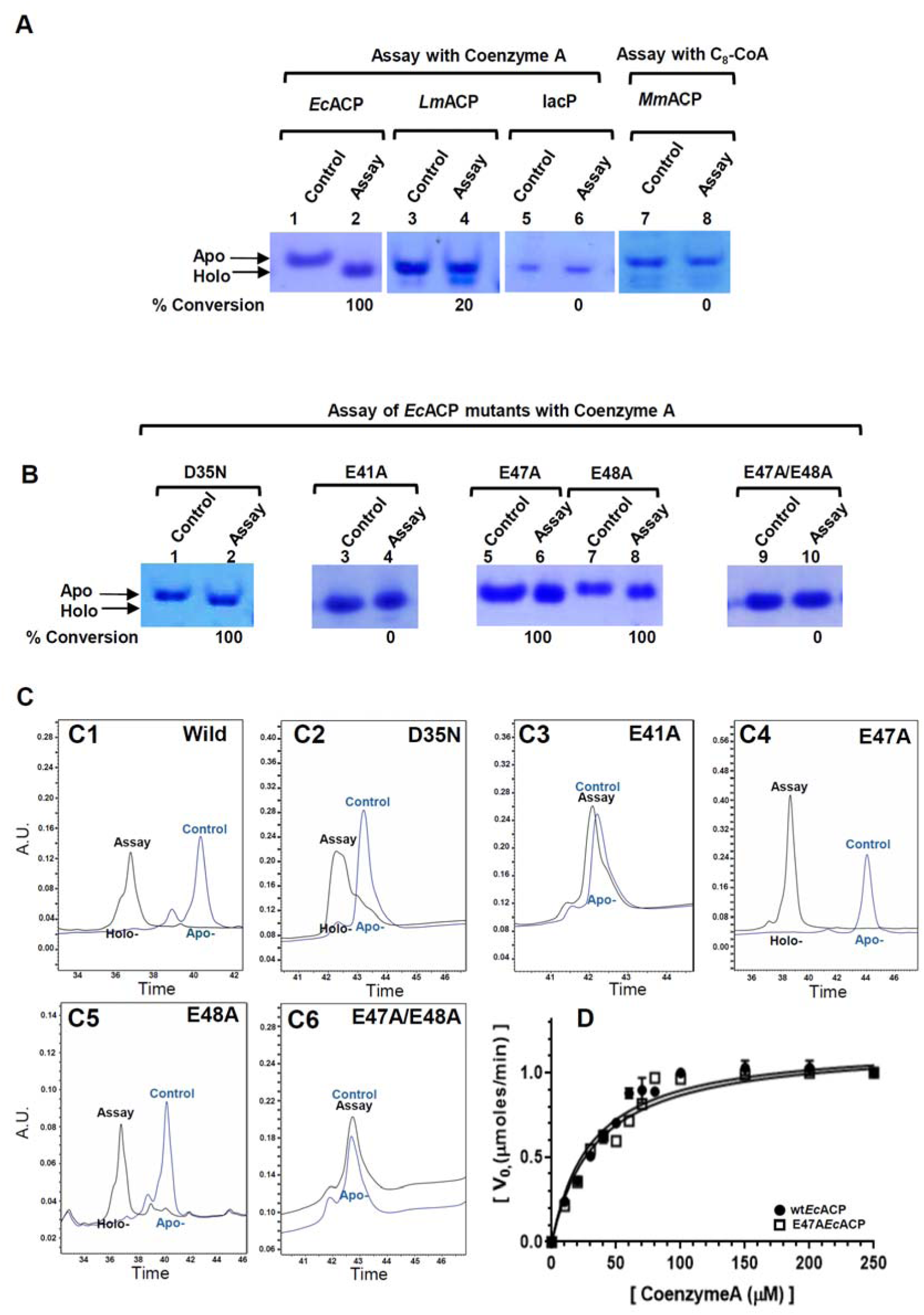
The extent of phosphopantetheinylation varies in different ACPs when expressed in *E. coli*. A 12% Native-PAGE gel displaying the bands for ACP before and after Sfp assay for A) *Ec*ACP (lanes 1 & 2), *Lm*ACP (lanes 3 & 4), IacP (lanes 5 & 6) and *Mm*ACP (lanes 7 & 8). B) Control and assay samples displaying the bands for *Ec*ACP mutants D35N (lanes 1 & 2), E41A (lanes 3 & 4), and E47A/E48A (lanes 5 & 6). Percent conversion of ACP after Sfp assay is listed at the bottom of the lane in each case, calculated using C_18_-reverse phase chromatography. C) C_18_-reverse phase chromatograms for the control-(blue) and Sfp assay-(black) samples for C1) wild type apo-*Ec*ACP, C2) D35N, C3) E41A, C4) E47A, C5) E48A, and C6) E47A/E48A*Ec*ACP. D) A plot of reaction velocity as a function of CoA concentration for *Ec*ACP and E47A mutant.

The apo-/holo-form of the expressed ACPs were further validated using ESI-MS spectra. Figure S1A displays the deconvoluted ESI-MS spectra for IacP, and Fig. S1C the charge envelope for *Lm*ACP. The molecular mass of IacP determined by ESI-MS corresponds to holo-ACP, while the mass of *Lm*ACP is equivalent to the apo-form. For comparison, ESI-MS spectra for apo-and holo-*Ec*ACP (Fig. S1E & F) were also recorded. Mass spectra was acquired for *Mm*ACP as well, and the determined isotopic mass was found to be higher than the molecular mass of holo-ACP (corresponding to C_18_-ACP). Previous studies with human mitochondrial ACP have also indicated that the protein is expressed primarily in the acylated form in *E. coli* (*14*). These results unequivocally show that IacP and *Mm*ACP are expressed as modified proteins, while *Lm*ACP is expressed in the apo-form in *E. coli*. *Ec*ACP is expressed as a partly modified protein.

### Charge neutralization mutations of *Ec*ACP prevent its *in vivo* modification by AcpS

Holo-acyl carrier protein synthase (AcpS) is a type I phosphopantetheinyl transferase (PPTase), endogenously present in *E. coli*. Due to its broad substrate specificity, AcpS can modify several recombinant acyl carrier proteins (ACPs), in addition to *Ec*ACP. AcpS is a homotrimer, consisting of 3 equal subunits. Each trimer can bind 3 ACP molecules. The crystal structure of *E. coli* AcpS trimer (UniProtKB P24224, colored orange) in complex with three holo-*Ec*ACP molecules (PDB 5VCB, colored blue) is shown in Fig. 3A. The interaction interface is formed by loop I-helix II of *Ec*ACP and helix I of AcpS. In Fig. 3B, the ACP-AcpS interaction interface is shown for chains A & S, respectively (PDB 5VCB). The charged residues present at the protein-protein interaction interface are shown in ball and stick, colored by hetero atom. Specifically, four arginines of AcpS (Arg 15, Arg 22, Arg 26 and Arg 30), and five carboxylates of *Ec*ACP, (Asp 35, Asp 38, Glu 41 and Glu 47 and Glu 48), are present in this region.

**Figure 3.**
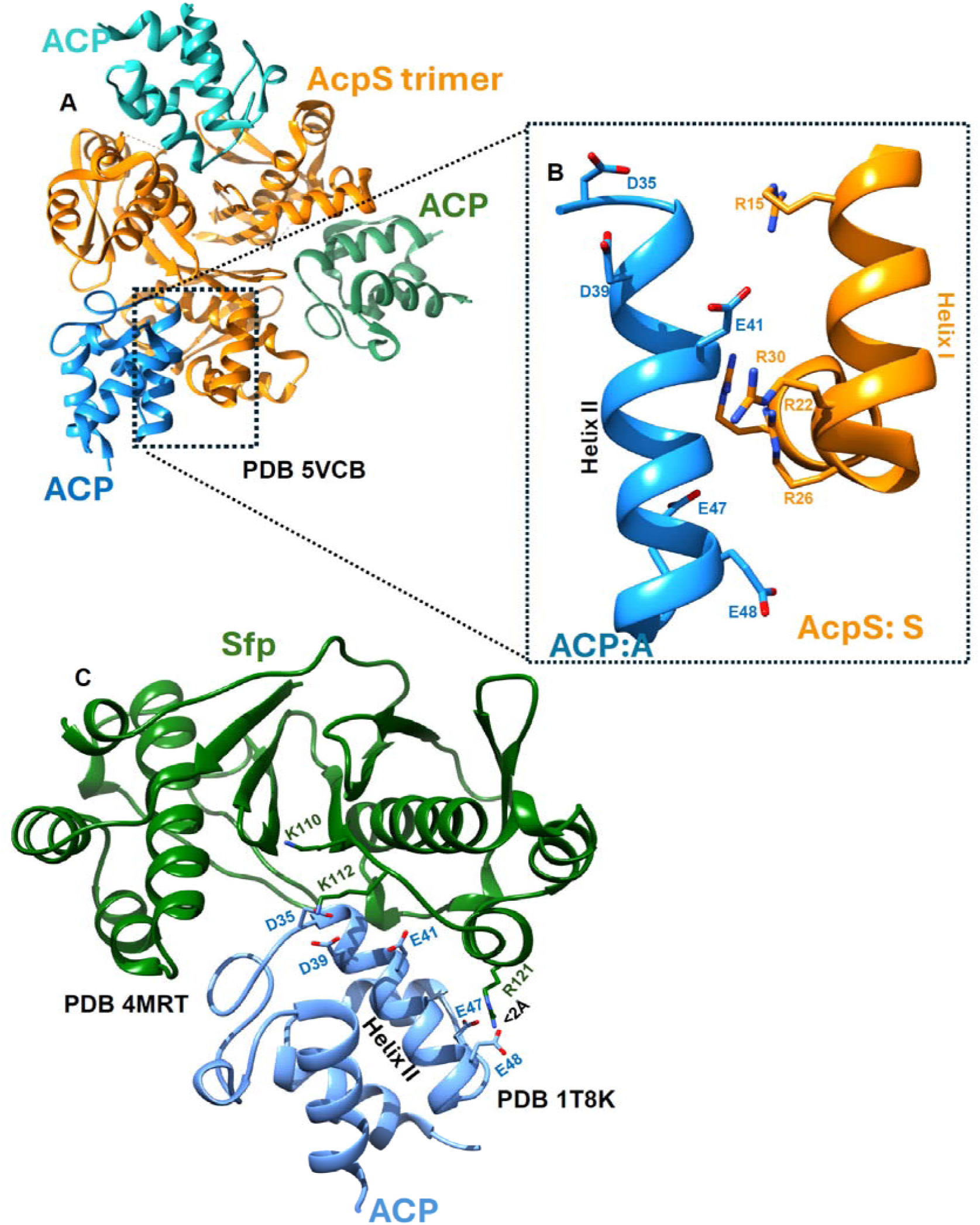
AcpS and Sfp differ in their interaction with ACP. A) A ribbon representation of *E. coli* Holo-acyl carrier protein synthase (AcpS trimer) in complex with three ACP molecules (PDB 5VCB). B) Electrostatic interactions between the helix II of *Ec*ACP and helix I of AcpS. The carboxylates present in the helix II of ACP and the arginines of AcpS are shown in ball and stick, colored based on hetero atom. C) A ribbon representation of *Bacillus subtilis* Sfp in complex with *Ec*ACP (PDB 1T8K). The figure was generated by docking the ACP structure (PDB 1T8K) on the PCP molecule in PDB 4MRT using the matchmaker option of Chimera(*25*).

Given the electrostatic nature of interaction between the two proteins, we hypothesized that charge neutralization mutations of *Ec*ACP could prevent productive ACP-AcpS complex formation, and thereby limit *in vivo* phosphopantetheinylation. The apo-mutants so expressed can be modified later using Sfp *in vitro*. To test this hypothesis, five *Ec*ACP mutants, D35N, E41A, E47A, E48A and a double mutant E47A/E48A were generated. Aspartate 35 was mutated to an Asn instead of Ala, since this variation exists naturally in *Leishmania major* ACP (*Lm*ACP), shown in Fig 1A (colored red). This conservative mutation has also been reported in a previous study (*26*).

Thus, *Ec*ACP and its mutants were expressed in *E. coli,* and purified using anion exchange chromatography. For comparison, *Lm*ACP was also expressed and purified. The fractions obtained after elution through a resource Q column were loaded on a 12% Native-PAGE. Wt*Ec*ACP (Fig. 4B), its mutants E47A (Fig. 4E) and E48A (Fig. 4F) displayed two distinct bands on native-PAGE, and the holo-ACP band could be seen migrating ahead of the apo-band. Chromatograms displaying two separate peaks for the apo-and the holo-forms of these proteins are shown in Fig. 4H-J. In contrast, *Lm*ACP (Fig. 4A), and the *Ec*ACP mutants D35N (Fig. 4C), E41A (Fig. 4D) and E47A/E48A (Fig. 4G) eluted as a single peak through the resource Q column (data not shown) and displayed a single band on native-PAGE in all the fractions suggesting one form of ACP.

**Figure 4.**
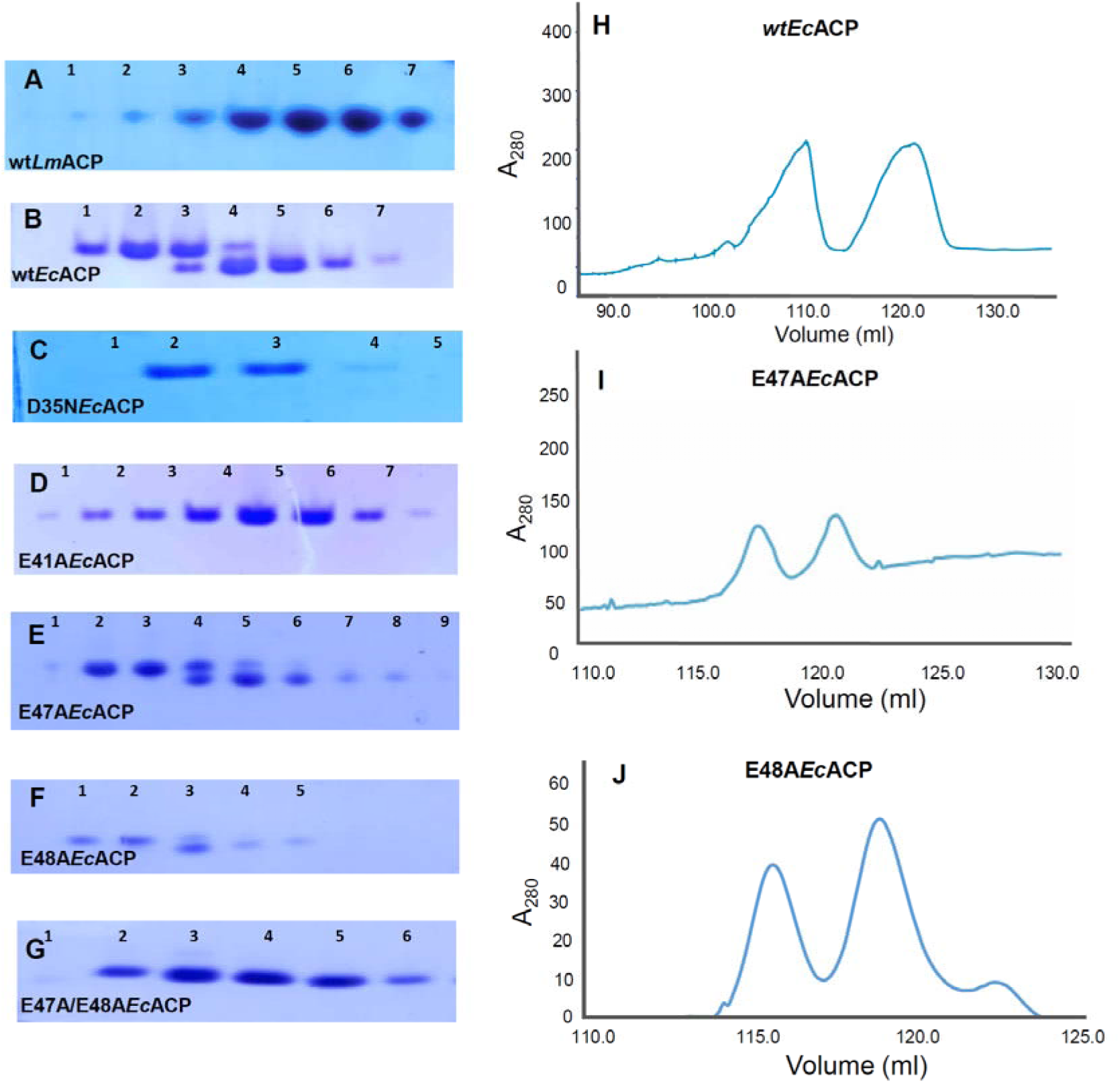
*In vivo* modification of wild-type ACPs and mutants expressed in *E. coli*. A 12% Native-PAGE gel displaying the fractions obtained after anion-exchange chromatography for A) *Lm*ACP, B) wt*Ec*ACP, *Ec*ACP mutants C) N35D, D) E41A, E) E47A, F) E48A, and G) E47A/E48A. Chromatogram obtained after elution of proteins through a Cytiva 6ml Resource Q column are shown for H) wt*Ec*ACP, I) E47A and J) E48A, displaying two separate peaks for apo-and holo-ACP.

To confirm the identity of the mutants, ESI-MS spectra were recorded for: *Lm*ACP after Sfp assay (Fig. S1D), *Ec*ACP mutants apo-E47A (Fig. S1G), holo-E47A (Fig. S1H), apo-E48A (Fig. S1I), holo-E48A (Fig. S1J) and E47A/E48A*Ec*ACP double mutant (Fig. S1K). The observed isotopic mass of *Lm*ACP after Sfp assay was consistent with the mass of the holo-form, and E47A/E48A*Ec*ACP (Fig. S1K) with the apo-form. In all cases, the ESI-MS determined masses were within ±2 Da of the expected value.

### Mutations induce small chemical shift perturbations in *Ec*ACP

To understand the effect of mutations on the *Ec*ACP structure, ^1^H^15^N HSQC spectra were acquired for apo-*Ec*ACP and its mutants. In Fig. S2 (panels B-E), ^1^H^15^N HSQC spectra are shown for the *Ec*ACP mutants (colored green) D35N (Fig. S2B), E41A (Fig. S2C), E47A (Fig. S2E) and E47A/E48A (Fig. S2F) overlaid on the wild type apo-*Ec*ACP spectrum (colored red). For comparison, ^1^H^15^N HSQC spectrum for S36A*Ec*ACP mutant (colored green) superimposed on the wild type apo-*Ec*ACP spectrum (colored red) is also shown in Fig S2A. All the proteins displayed good spectral dispersion, indicative of a well-structured protein. Minor chemical shift perturbations were observed at the amides of a few residues in D35N, S36A and E41A mutants, suggesting insignificant structural changes upon mutagenesis. However, E47A (Fig S2E), and E47A/E48A (S2F) displayed slightly larger perturbations, probably due to the loss of E47 sidechain-Ile 54 backbone hydrogen bond, disclosed by the 1.3 ppm upfield change in the chemical shift of Ile 54 amide in the two mutants (Fig. S2D).

### Sfp induces chemical shift perturbations in the helix I, loop I and helix II amides of *Ec*ACP

Since the crystal structure of ACP-Sfp complex has not been solved, the structure of Sfp-PCP complex (PDB 4MRT) solved at 2Å resolution was used to gain insights into the Sfp-carrier protein interactions. Peptidyl carrier proteins (PCP) are quite different in sequence compared to FAS ACP, but structurally they are similar. Therefore, *Ec*ACP (PDB 1T8K) was docked onto the PCP structure (PDB 4MRT:C), using the matchmaker option of Chimera(*25*). Upon careful examination of the Sfp-ACP docked complex (based on PDB 4MRT) (Fig. 3C), several key residues in the helix I of Sfp (Lys 110, Lys 112 and Arg 121) were found to be in the proximity of loop I and helix II residues (Asp 35, Asp 38, Glu 41 and Glu 47) of *Ec*ACP.

To further probe the *Ec*ACP-Sfp interaction in solution, NMR titration studies were carried out using ^15^N labeled S36A*Ec*ACP and unlabeled Sfp. The S36A mutant was specifically chosen for this study to prevent unwanted modification of ACP-Ser 36 in the presence of excess of Sfp. The *Ec*ACP spectrum was assigned based on the BMRB ID 27872.

Upon addition of increasing concentrations of Sfp, several amides in the ^1^H^15^N HSQC-TROSY spectrum of S36A*Ec*ACP displayed line broadening, an indicator of intermediate exchange, and enzyme-substrate complex formation. In the overlaid spectra shown in Fig. 5A, peaks corresponding to the unbound S36A*Ec*ACP spectrum are colored red-yellow, while the peaks for ACP bound to Sfp at 1:2 molar ratio are colored green. Due to line broadening, several ACP peaks in the Sfp bound state (green) were not visible in the spectrum. Figure 5B shows a plot of the relative peak intensity change (I/I_0_) as a function of residue number for ^1^H^15^N labeled S36A*Ec*ACP at 1:2 ACP:Sfp molar ratio. A dotted line in the figure represents ±1SD and has been used as an indicator of significant change. The residues displaying intensity change greater than ±1SD were Ile 10, Gly 12, Gln 14, Val 17-Gln 19, Thr 23, Ala 26, Ser 27-Asp 35, Asp 38, Val 40-Met 44 and Glu 47. These amides have been colored pink in the ribbon representation of *Ec*ACP docked on the Sfp molecule, shown in Fig. 5C. Glutamate 4, Glu 6, Glu 22, Asn 24, Leu 15 and Thr 39 displayed overlap with other residues in the spectrum and therefore were not included in the analysis. These positions have been shown as an asterisk along x-axis in the plot. Notably, most of the residues displaying line broadening were hydrophobic, and the interaction surface encompassed four carboxylates as well, namely Asp 35, Asp 39, Glu 41 and Glu 47. From the titration it appears that Sfp interacts primarily with the loop I and helix II residues of *Ec*ACP, consistent with the Sfp-PCP crystal structure. The perturbations observed in helix I could be due to the strain induced as a result of Sfp binding.

**Figure 5.**
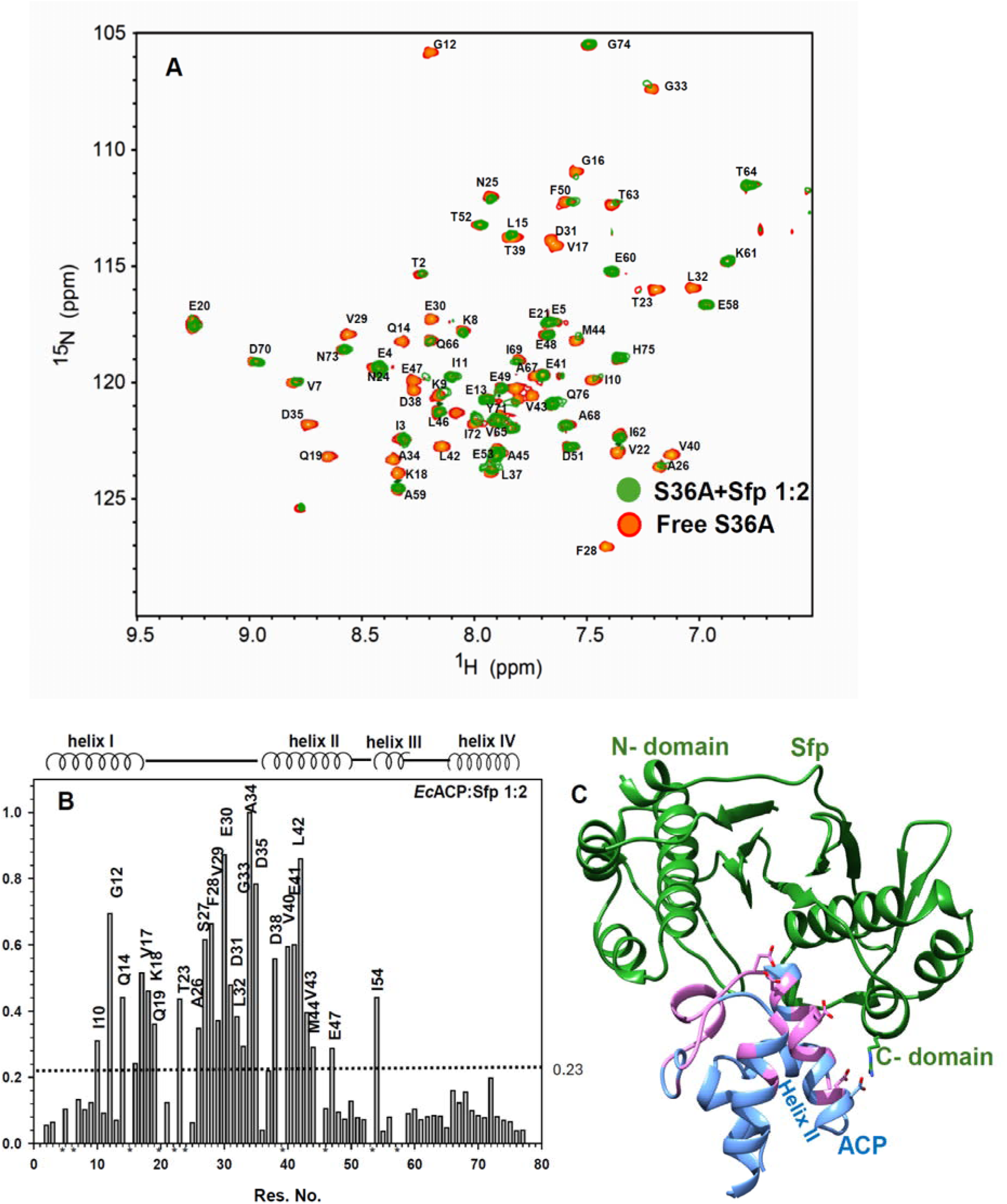
Chemical shift perturbations of *Ec*ACP upon interaction with Sfp. A) ^1^H^15^N TROSY-HSQC spectrum for S36A*Ec*ACP in complex with Sfp in 1:2 molar ratio (peaks colored green) overlaid on the S36A*Ec*ACP spectrum (peaks colored yellow-red). The figure was prepared using Sparky (*19*). B) A plot of the relative intensity change as a function of residue number for S36A*Ec*ACP amides upon interaction with Sfp (in 1:2 molar ratio). A discontinuous horizontal line in the plot indicates one standard deviation. Some of the amides that displayed overlap in the spectrum were not included in the analysis and have been indicated by an astersk * along x-axis. C) A ribbon representation of *Ec*ACP (PDB 1T8K, colored blue) docked on the PCP molecule in complex with *Bacillus subtilis* Sfp (PDB 4MRT, colored green). The S36A*Ec*ACP amides displaying significant change in peak intensity upon Sfp interaction are colored pink.

### D35N*Ec*ACP was successfully modified with Sfp *in vitro*

I*n vitro* phosphopantetheinylation of the *Ec*ACP mutants D35N, E41A, and the double mutant E47A/E48A*Ec*ACP was carried out using Sfp *in vitro*. For comparison, Sfp assays were also performed using apo-E47A, and apo-E48A mutants as substrates. Each mutant was incubated under two different conditions, 1) without Sfp (used as control), and 2) with Sfp (assay), in the presence of CoA. After the Sfp assay, both control and assay samples were loaded on a 12% Native-PAGE, to monitor the extent of enzymatic conversion. Assay samples for D35N (Fig 2B, lane #2), apo-E47A (lane #6), and apo-E48A (lane #8) *Ec*ACP mutants displayed full conversion to holo-ACP, disclosed by the forward migration of the assay band. However, the assay samples for E41A (lane #4), and the double mutant E47A/E48A*Ec*ACP (lane #10) did not show any change in migration compared to the control samples (lanes #3 & #9), suggesting that Sfp was unable to phosphopantetheinylate E41A and E47A/E48A *Ec*ACP mutants.

HPLC C_18_ reverse phase chromatography was used to quantitatively measure the extent of conversion in each case. Apo-*Ec*ACP (Fig. 2C1), its mutants D35N (Fig. 2C2), apo-E47A (Fig. 2C4), and apo-E48A*Ec*ACP (Fig. 2C5) displayed 100% conversion into the holo-form in the presence of Sfp *in vitro,* disclosed by the change in the position of the assay peak (colored black) compared to control (apo-, colored blue). However, the two mutants, E41A (Fig. 2C3) and E47A/E48A (Fig. 2C6) did not show any change in the elution time of the assay peak compared to control, and the C_18_-reverse phase chromatogram profiles were identical for the control and assay samples, suggesting absence of any phosphopantetheinylation.

Kinetic parameters were also determined for Sfp using wt*Ec*ACP and E47A mutant as substrates. K_M_ and K_cat_ values were comparable for the two proteins, shown in Fig. 2D and Table-1. The data indicates that the loss of Glu 47 CO-Ile 54 hydrogen bond in the E47A mutant does not alter Sfp activity. Thus, the inability of Sfp to convert E47A/E48A*Ec*ACP double mutant (that also lacks the hydrogen bond) into holo-ACP cannot be attributed to structural alteration.

**Table 1.** Kinetic Constants for Sfp *(B. subtilis*) Using wt*Ec*ACP and E47A*Ec*ACP as Substrates.

| ACP | $K_M$ (μM) | $K_{cat}$ ($\text{min}^{-1}$) | $V_{max}$ ($\mu\text{Mmin}^{-1}$) | $K_{cat}/K_M$ ($\text{min}^{-1} \text{M}^{-1}$) |
| --- | --- | --- | --- | --- |
| wt*Ec*ACP | $31.19 \pm 3.6$ | $0.235 \pm 0.01$ | $1.175 \pm 0.03$ | 7534 |
| E47A*Ec*ACP | $34.62 \pm 3.7$ | $0.234 \pm 0.02$ | $1.168 \pm 0.03$ | 6759 |

Overall, of the five mutations introduced on the *Ec*ACP surface, three expressed as apo-ACP in *E. coli*. Of the three apo-ACP mutants, only D35N was successfully converted into holo-ACP using Sfp *in vitro*.

### D35N mutants of IacP and *Mm*ACP expressed as apo-ACP in *E. coli* and were readily modified by Sfp *in vitro*

The mutagenesis approach was tested on two remarkably distinct ACPs that express predominantly as modified proteins in *E. coli*; *Salmonella* Typhimurium invasion acyl carrier protein (IacP) and *Mus musculus* mitochondrial ACP (*Mm*ACP). D35NIacP and D35N*Mm*ACP, both expressed in only one form in *E. coli,* inferred from the observation of a single band on Native-PAGE. ESI-MS spectra for D35NIacP also suggests that the protein is expressed in its apo-form, shown in Fig. S2B.

Next, Sfp assays were performed with the D35N mutants of IacP and *Mm*ACP using different acyl-CoA as cosubstrates. Sfp assays were carried out with apo-*Ec*ACP as well (for comparison) In Fig. 6A, lanes #2-5 display the assays performed using apo-*Ec*ACP, lanes #7-10 using D35NIacP, and lanes #12-15 with D35N*Mm*ACP as substrates, using CoA, C_8_-CoA, C_10_-CoA and C_12_-CoA, respectively. The control samples (incubated without Sfp) are shown in lanes #1, #6 and #11 for apo-*Ec*ACP, D35NIacP and D35N*Mm*ACP, respectively. Sfp successfully converted apo-*Ec*ACP as well as the D35NIacP and D35N*Mm*ACP mutants into the holo-/acylated forms, disclosed by the faster migration of the assay bands compared to control. For some reason, *Mm*ACP displayed very less conversion with coenzyme-A, but could be fully modified using various acyl-CoAs as substrates.

**Figure 6.**
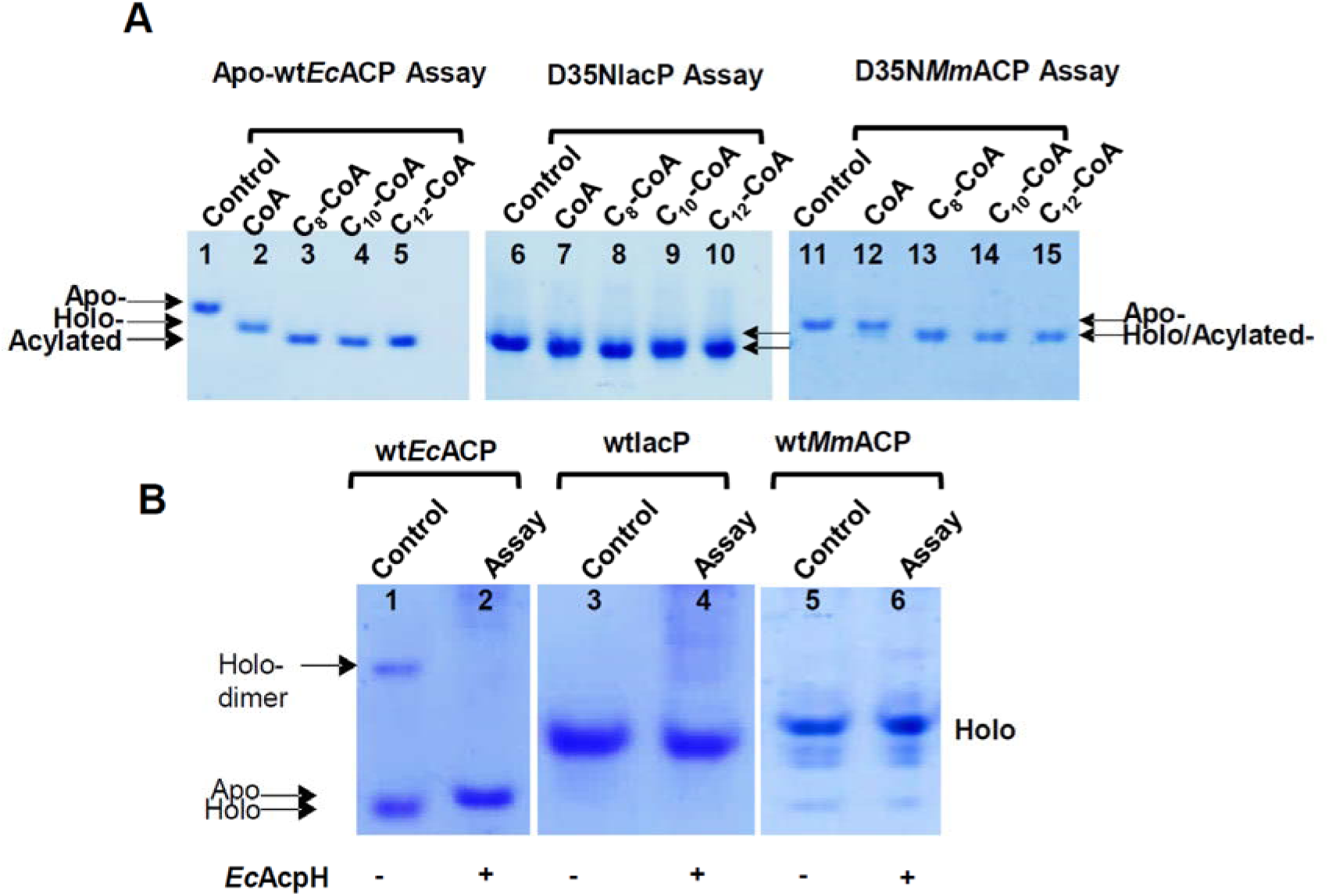
*In vitro* activity of Sfp and AcpH towards various ACPs. A) A 12% Native-PAGE gel displaying the bands for A1) apo-wt*Ec*ACP (lane #1-5), A2) D35NIacP (lane # 6-10) and A3) D35N*Mm*ACP (lane #11-15). In each case, the first lane displays the control sample (without Sfp), and the other four lanes display the Sfp assay carried out in presence of different CoA. The acyl-CoA used for a given reaction is mentioned at the top of each lane. An increase in the migration of the ACP band indicates conversion by Sfp. B) A 12% Native-PAGE gel displaying the conversion of holo-wt*Ec*ACP, holo-wtIacP and holo-wt*Mm*ACP into apo-ACP using AcpH. A noticeable decrease in the migration of the ACP band in the assay lane compared to control indicates conversion from holo-to apo-ACP.

The results were also verified using NMR. ^1^H^15^N HSQC spectra were acquired for the apo-and the acylated forms of the two D35N mutant proteins. Figure S3A displays the spectrum for C_4_-D35N*Mm*ACP (colored green), overlayed on the apo-35N*Mm*ACP spectrum (colored red). In Fig. S3B, C_2_-IacP spectrum (colored green) has been superimposed on the apo-D35NIacP spectrum (colored red). Residues of IacP were assigned based on previously published assignments (*27*). Changes in the position of some of the amide peaks, in particular Ser 36 and its neighboring residues confirm conversion of apo-IacP into C_2_-IacP.

### *E. coli* ACP hydrolase (AcpH) fails to revert modified-IacP and *Mm*ACP into the apo-form

An important question remains to be addressed; why certain ACPs such as IacP and *Mm*ACP are expressed predominantly in the holo-/acylated form in *E. coli,* while others like *Ec*ACP, and *Pf*ACP (*P. falciparum* ACP) are only partially modified? To find an answer to this question, *in vitro* assays were performed using *E. coli* acyl carrier protein hydrolase (AcpH), a well-known enzyme that can release the phosphopantetheine arm (4’-PP) from holo-ACP. AcpH assays were performed using three different substrates: holo-*Ec*ACP, IacP and *Mm*ACP, shown in Fig. 6B. Since IacP and *Mm*ACP express as modified proteins in *E. coli*, the nickel affinity purified protein was used in this assay. Holo-*Ec*ACP (lane #1) could be fully converted into the apo-form (lane #2) by AcpH, reported by the slow migration of the protein band in the assay sample compared to control (without AcpH). However, the assays performed with IacP (lane #4) and *Mm*ACP (lane #6) did not show any change in migration compared to the control bands (lanes #3 and #5), suggesting that AcpH is unable to convert phosphopantetheinylated-IacP and *Mm*ACP into the apo-form.

## DISCUSSION

Overexpression in bacteria is the standard protocol to obtain milligram quantities of a carrier protein. However, undesired modification is a frequently encountered problem, arising due to the recognition of non-native ACPs as a substrate by AcpS, a 4’-Phosphopantetheinyl transferase native to *E. coli*. Thus, additional purification steps are required, resulting in low yields of the apo-protein. The problem is exacerbated if the ACP is expressed as a fully modified protein. To circumvent this problem, we propose a strategy that specifically targets sites on ACP that interact with AcpS, without altering its interaction with Sfp. Since *E. coli* ACP-AcpS interactions are primarily electrostatic, while the PCP-Sfp interactions are hydrophobic, differences in the interaction of the two enzymes were exploited to specifically block ACP modification by AcpS. Thus, carboxylates in the loop I-helix II of ACP that interact with the arginines of AcpS were identified as hot spots for mutation. Aspartate 38 was not mutated, since it is necessary for Sfp activity (*28*). Charge neutralization mutations introduced at three different positions in *Ec*ACP-35, 41, and 47/48, resulted in the expression of apo-ACP in *E. coli*. However, only the D35N mutant could be successfully converted into holo-/acyl-ACP by Sfp *in vitro*. A previous study with Sfp-R-4-4, a phage display optimized mutant has shown that the enzyme can phosphopantetheinylate E41A and several other *Ec*ACP mutants due to its extremely broad substrate specificity (*29*). Possibly, Sfp-R-4-4 if available could convert all three *Ec*ACP mutants into the holo-/acylated-form. Since in our studies, D35N*Ec*ACP was the only mutant that successfully converted into the holo-and acylated forms with wild-type Sfp, the mutation emerged as an attractive strategy to express unmodified ACPs in *E. coli*. As proof of concept, D35N mutation was introduced in two other rather challenging systems, IacP (*Salmonella* Typhimurium invasion acyl carrier protein) and *Mm*ACP (*Mus musculus* mitochondrial ACP), that express primarily as holo-/ acylated-proteins in *E. coli* (*14*). Interestingly, the D35N mutants of *Mm*ACP and IacP also expressed in the apo-form in *E. coli* and smoothly converted into the holo-/acylated-form with Sfp, establishing D35N mutagenesis as a viable strategy to express apo-proteins across all type II ACPs.

Though some of the *Ec*ACP charge neutralization mutants were not modified by Sfp, the results indirectly pointed towards the role of charged residues in Sfp interaction. All the studies published to date emphasize the importance of hydrophobic interactions in Sfp recognition. For instance, the crystal structure of Sfp-PCP complex highlights the indispensable role of hydrophobic interactions, apart from an electrostatic interaction involving Asp 38 of ACP (Tufar et al., 2014) (*28*). A recent mutagenesis study uncovered the role of hydrophobic residues in making ACP/PCP Sfp compatible(*30*). Similarly, a phage display study identified A1 peptide ‘GD<u>S</u>LDMLEWSLM’ as an AcpS substrate, while S6 peptide ‘GD<u>S</u>LSWLLRLLN’ was found suitable for Sfp (*31*). Sfp was unable to phosphopanthetheinylate A1 peptide due to the presence of a Glu at position 8, instead of a leucine. Adding to the current understanding of Sfp-ACP interaction, our studies show that electrostatic interactions also play a role in productive Sfp-ACP interaction. Sfp was able to fully modify the charge neutralization single mutants E47A, and E48A*Ec*ACP. However, when both the charges were removed in the double mutant E47A/E48A, no phosphopantetheinylation was observed. NMR titration studies using ^1^H^15^N E47A/E48A*Ec*ACP and Sfp display line broadening, suggestive of enzyme-substrate complex formation (data not shown), but it never leads to the modification of ACP. Thus, a charged residue at position 47 or 48 in ACP is necessary for its productive interaction with Sfp. These observations are in compliance with the amino acid sequence of PCPs’ (natural substrates of Sfp), that have a conserved polar/charged residue at positions 47 and 48 (*29*).

Furthermore, our studies provide explanation for the expression of fully modified ACPs in *E. coli* (*14*). *E. coli* Acyl carrier protein hydrolase (AcpH) failed to convert modified-IacP and *Mm*ACP into apo-ACP, though it successfully released 4’-PP moieties from holo-*Ec*ACP. These results indicate that modified-IacP and *Mm*ACP are probably not suitable AcpH substrates. Taken together, it emerges that the extent of modification of an overexpressed ACP is dependent on the catalytic activity of two *E. coli* enzymes, AcpS and AcpH, that display antagonistic activity. The substrate of one enzyme is the product of the other, and *vice versa*. In cases where AcpH is unable to recognize ACP as a substrate (due to differences in sequence with its cognate substrate), fully modified ACPs are expressed, *e.g.* mitochondrial human ACP. When a non-native ACP does not meet the substrate requirements of AcpS, the latter is expressed as an apo-protein, *e.g*. *Leishmania* ACP. However, when both AcpS and AcpH are functional, a partly modified ACP population is obtained, as observed in the case of *E. coli, P. falciparum, M. tuberculosis etc*. Thus, by modulating AcpS or AcpH activity, it is possible to express 100% apo-or holo-ACP in *E. coli*.

### Accession IDs

- UniProtKB P0CL00
- UniProtKB Q569N0
- UniProtKB P21515
- UniProtKB E9AD06
- UniProtKB P0A6A8
- UniProtKB P39135

## Author Contributions

C.D., D.S., R.P., and S.V. contributed equally to the work. The names are listed in alphabetical order.

## Data Availability Statements

The data supporting the findings of this study are available within the paper and in Supporting Information. Additional files are available from the corresponding author on reasonable request.

## ACKNOWLEDGMENTS

The *L. major* genome was obtained through BEI Resources, NIAID, NIH: genomic DNA from L. major strain NIH SD (MHOM/SN/74/SD), NR-48764. The Genomic DNA of *Salmonella enterica* subsp. *enterica*, 2004 Pennsylvania Tomato Outbreak, Serovar Typhimurium was obtained through BEI Resources, NIAID, NIH: Isolate 1, NR 4614. Authors thank the Dept of Biotechnology, Govt. of India for financial and infrastructure support. Fellowship from the Lady Tata Memorial Trust, India to Chetna Dhembla is gratefully acknowledged. Authors thank Ms. Shanta Sen and Ms. Richa Arya, Mass-Spec Facility NII, for help with the ESI-MS data.

## ABBREVIATIONS

ACP: Acyl carrier protein
*Ec*ACP: *E. coli* acyl carrier protein
*Mm*ACP: *Mus musculus* acyl carrier protein
IacP: Salmonella Typhimurium invasion acyl carrier protein
*Lm*ACP: *Leishmania major* ACP
*mt*ACP: Mitochondrial ACP
FAS: fatty acid synthase
CoA: coenzyme A
PPT: phosphopantetheinyl transferase
Sfp: *Bacillus subtilis* type II phosphopantetheinyl transferase
AcpS: Holo-acyl carrier protein synthase
AcpH: Holo-acyl carrier protein phosphodiesterase

