## Supplementary Figures for "A single mutation in the ‘DSL’ motif of acyl carrier protein can prevent its *in vivo* modification by *E. coli* Holo-acyl carrier protein synthase (AcpS)"

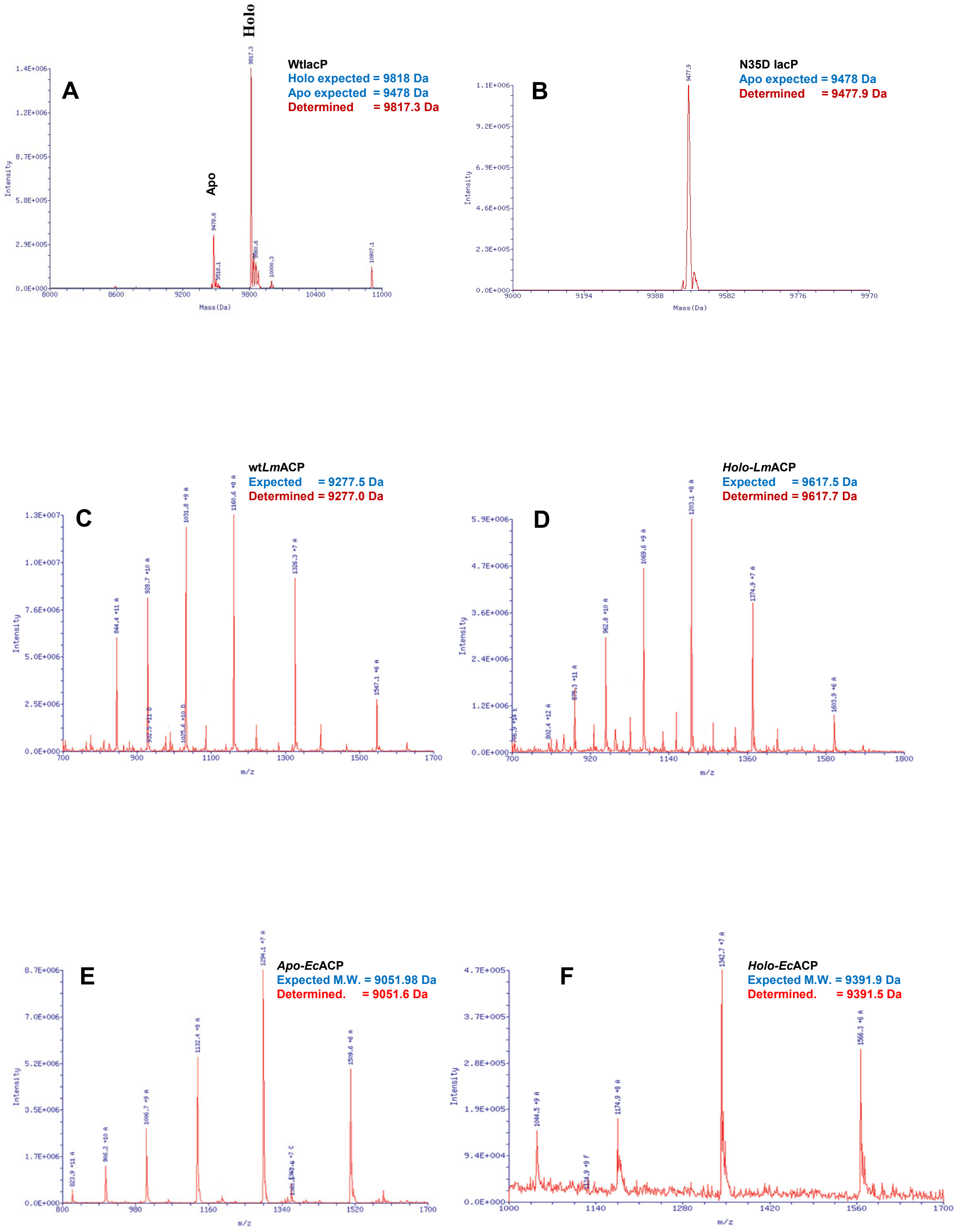
A
Holo
Apo
WtlacP
Holo expected = 9818 Da
Apo expected = 9478 Da
Determined = 9817.3 Da
Intensity
Mass(Da)
B
N35D lacP
Apo expected = 9478 Da
Determined = 9477.9 Da
Intensity
Mass(Da)
C
wtLmACP
Expected = 9277.5 Da
Determined = 9277.0 Da
Intensity
m/z
D
Holo-LmACP
Expected = 9617.5 Da
Determined = 9617.7 Da
Intensity
m/z
E
Apo-EcACP
Expected M.W. = 9051.98 Da
Determined. = 9051.6 Da
Intensity
m/z
F
Holo-EcACP
Expected M.W. = 9391.9 Da
Determined. = 9391.5 Da
Intensity
m/z

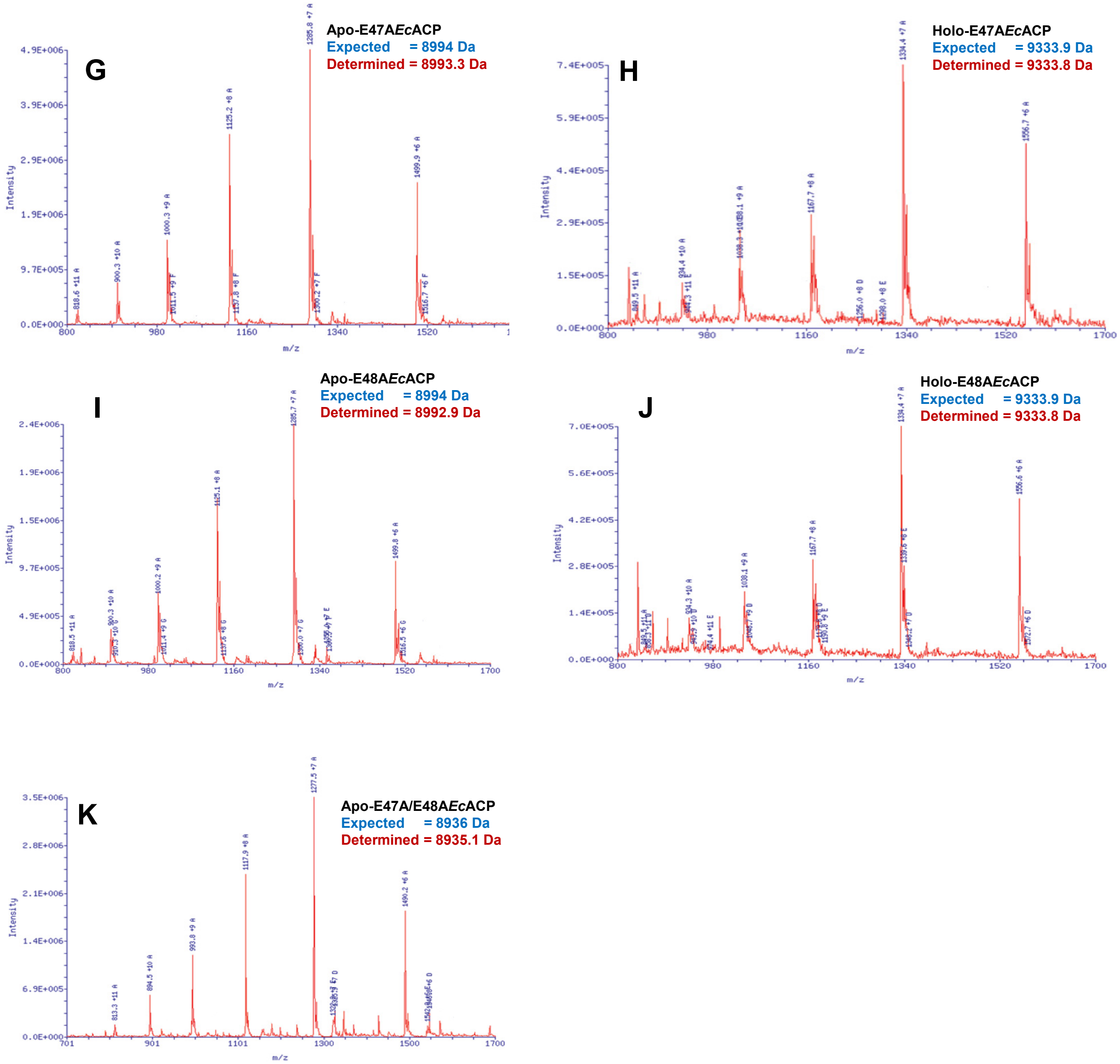


**Figure S1** ESI-MS spectra for various ACPs used in the study. A-B) Deconvoluted ESI-MS spectra for wtIacP and D35NIacP. Isotopic envelopes for C) *Lm*ACP, D) holo-*Lm*ACP (after Sfp assay), E) apo-wt*Ec*ACP, F) holo-wt*Ec*ACP, G) apoE47A, H) Holo-E47A, I) apo-E48A, J) holo-E48A, K) E47A/E48A*Ec*ACP.

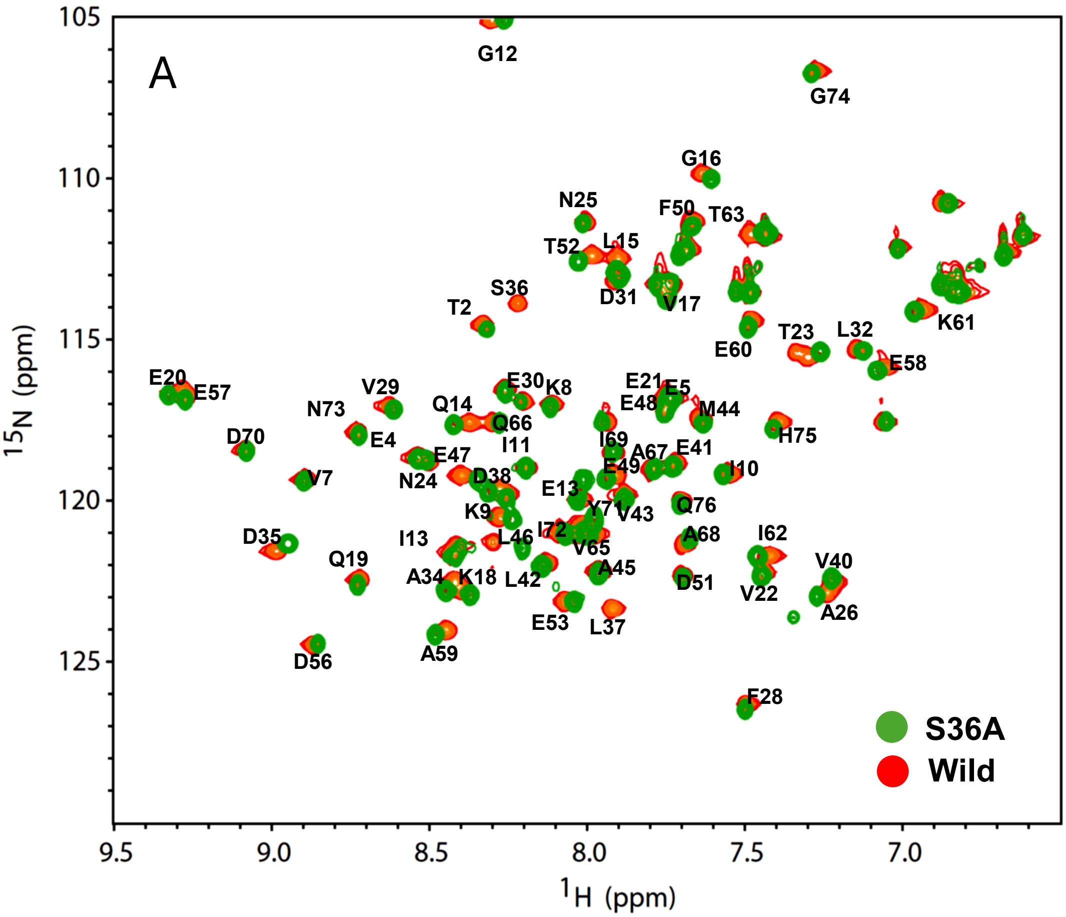
A
S36A
Wild
15N (ppm)
1H (ppm)
G12
G74
G16
N25
F50
T63
T52
L15
S36
T2
D31
V17
K61
T23
L32
E60
E58
E20
E57
V29
E30
K8
E21
E5
N73
Q14
E48
M44
D70
E4
Q66
I69
H75
E47
I11
A67
E41
V7
N24
D38
E49
I10
E13
K9
Y71
V43
Q76
D35
I13
L46
I72
A68
I62
V65
Q19
A34
K18
L42
A45
D51
V40
V22
E53
L37
A26
D56
A59
F28

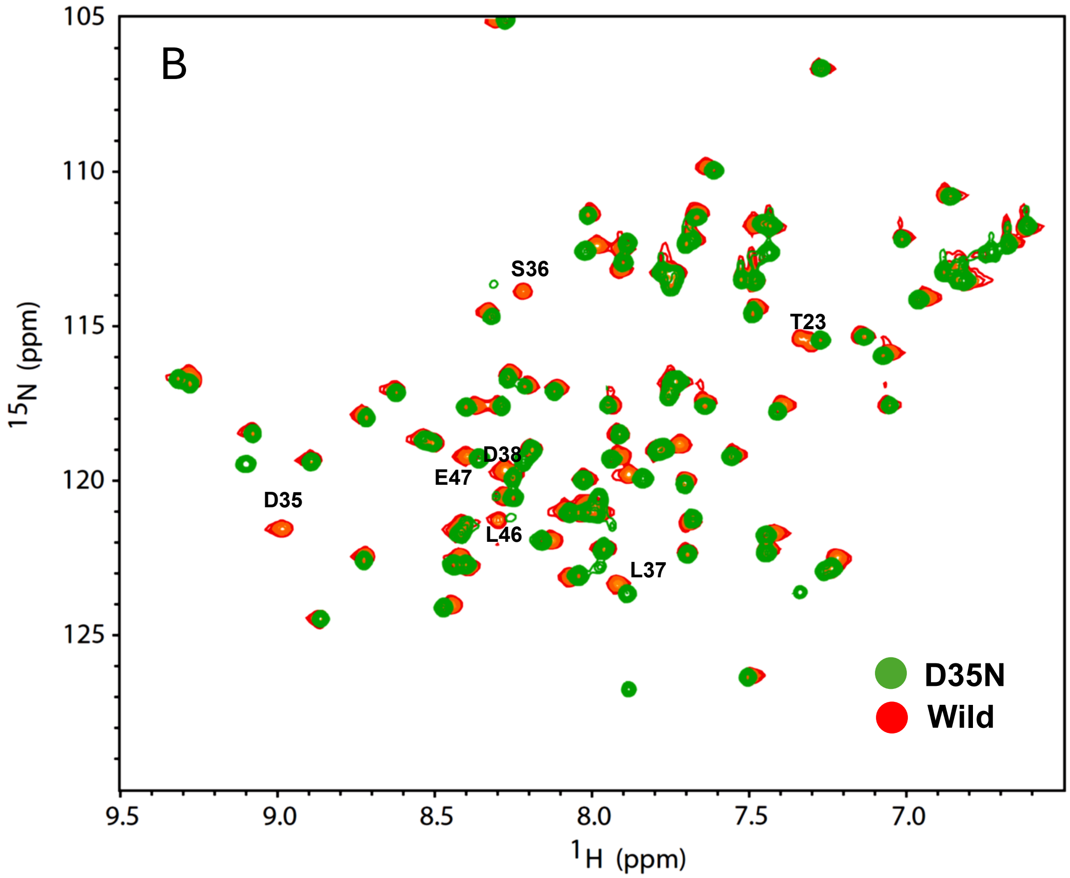
B
D35N
Wild
15N (ppm)
1H (ppm)
S36
T23
D38
E47
D35
L46
L37

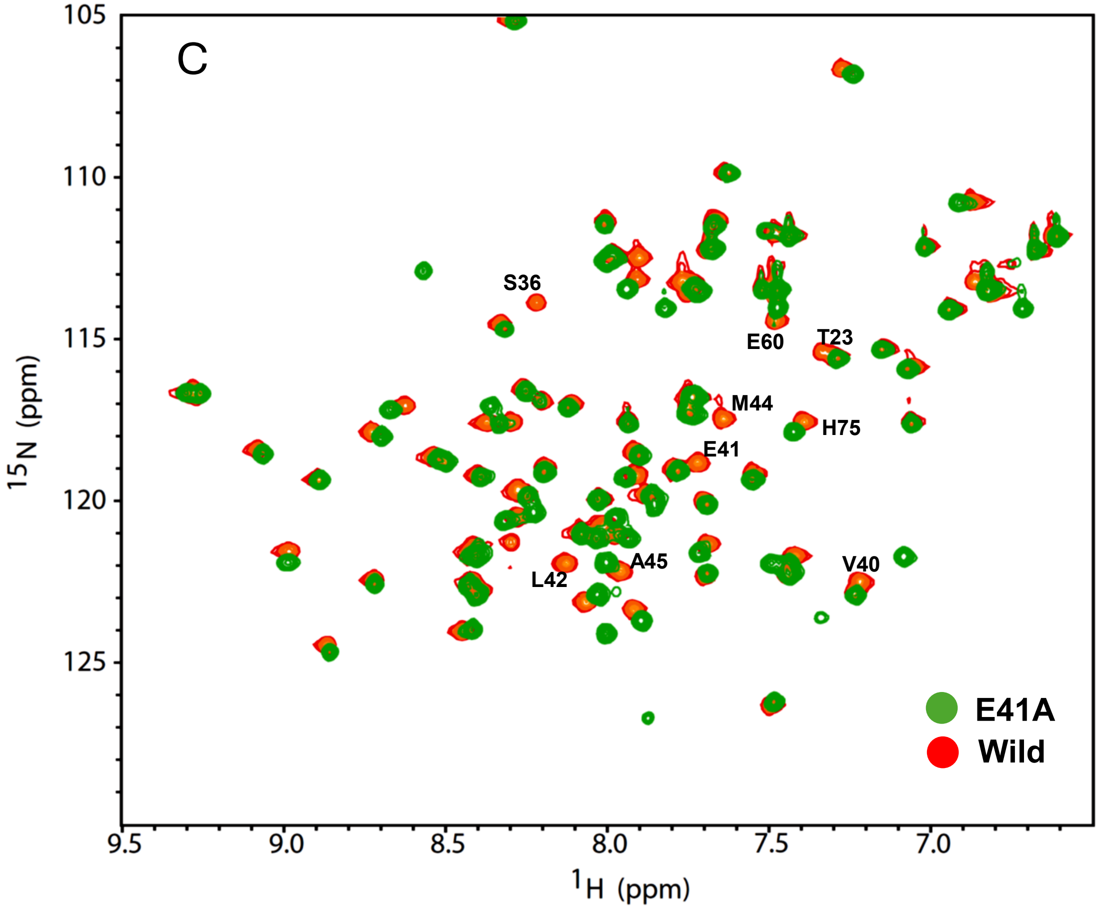
C
S36
E60
T23
M44
H75
E41
A45
L42
V40
E41A
Wild
105
110
115
120
125
15N (ppm)
9.5
9.0
8.5
8.0
7.5
7.0
1H (ppm)

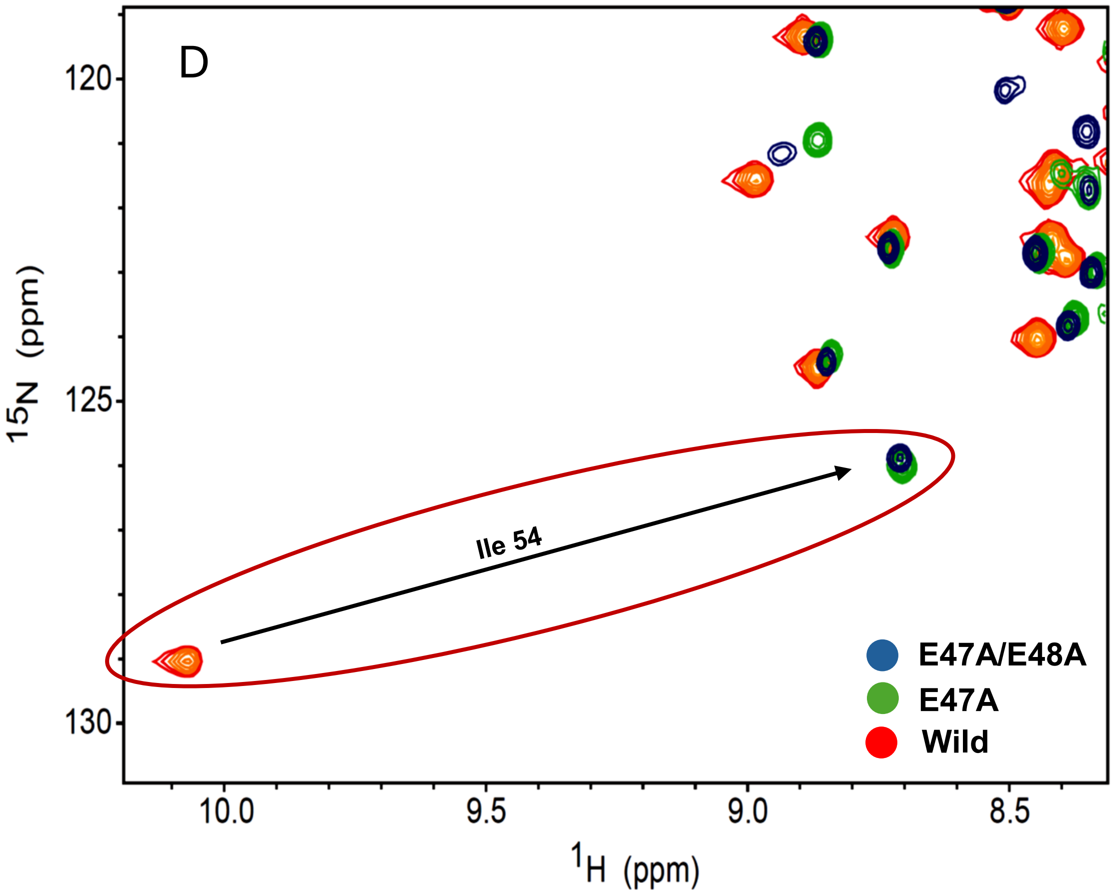
D
Ile 54
E47A/E48A
E47A
Wild
120
125
130
15N (ppm)
10.0
9.5
9.0
8.5
1H (ppm)

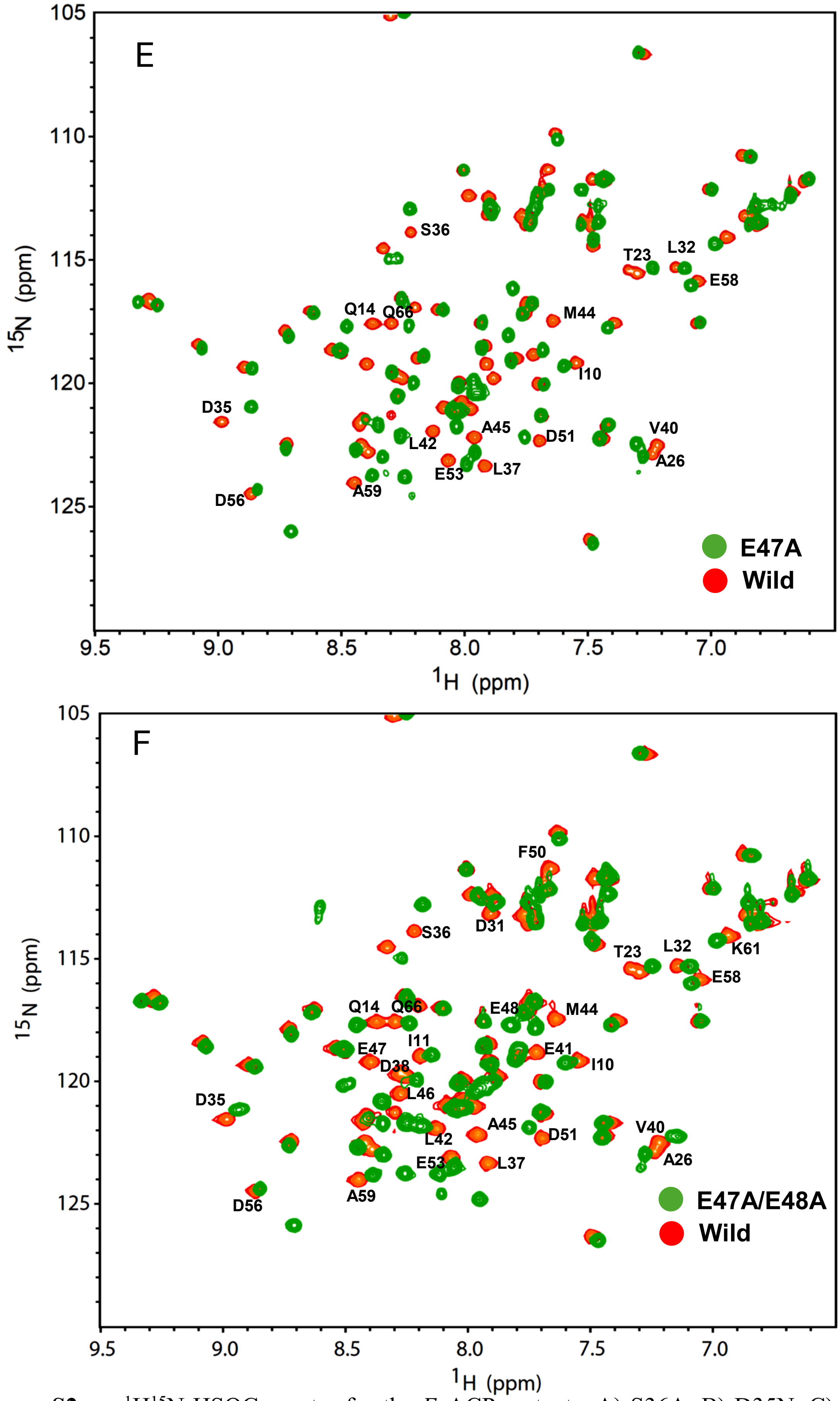


**Figure S2** $^{1}H^{15}N$ HSQC spectra for the *Ec*ACP mutants. A) S36A, B) D35N, C) E41A D) Glu 47-Ile 54 hydrogen bond in E47A and E47A/E48A, E) E47A, and F) E47A/E48A*Ec*ACP spectra (colored green) overlaid on the apo-wt*Ec*ACP spectra (colored red). Some of the residues that display chemical shift changes have been marked. The figures were prepared using Sparky *(19)*.

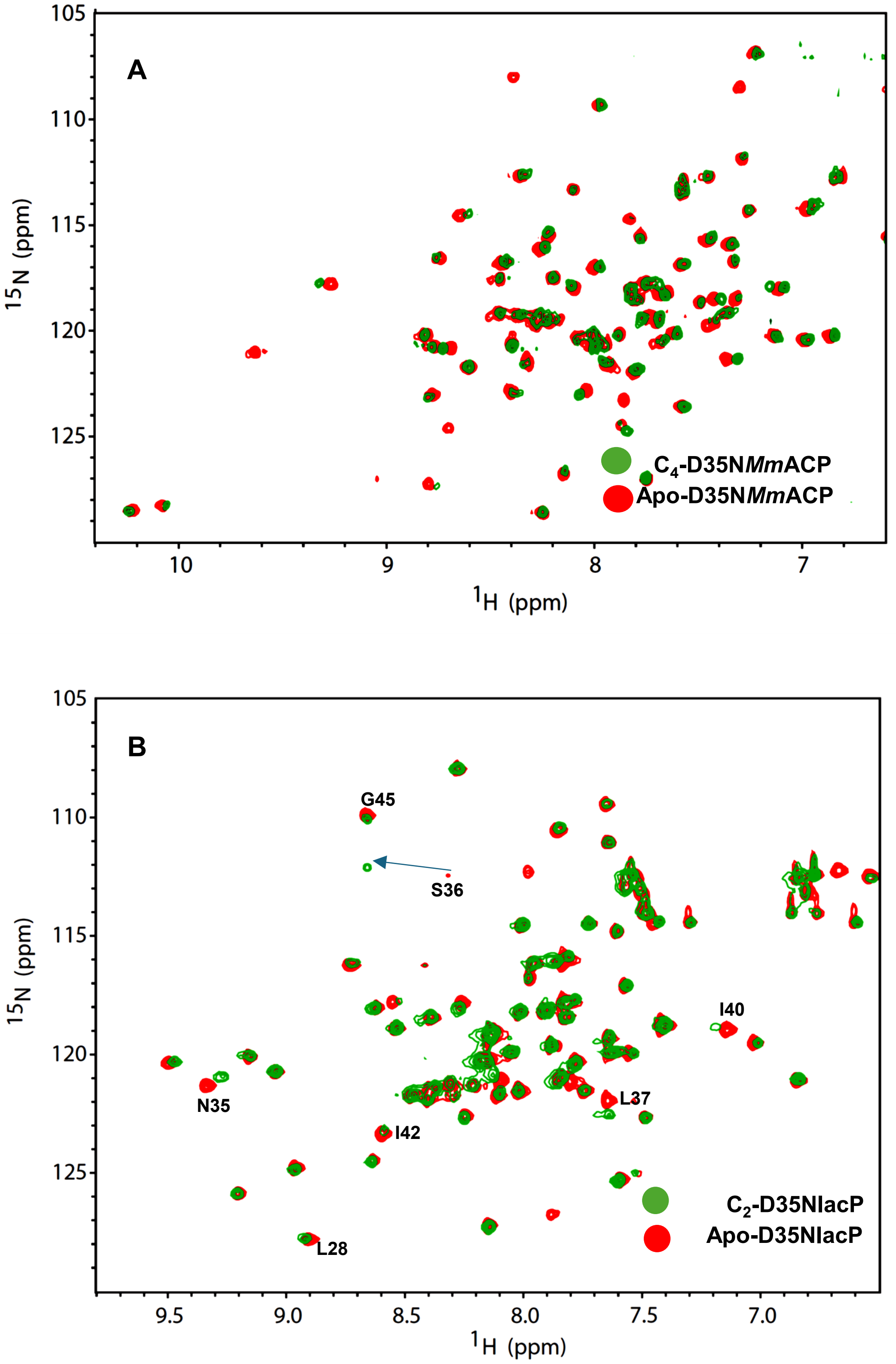


**Figure S3** $^{1}H^{15}N$ HSQC spectra for D35N*Mm*ACP and D35NIacP. A) $C_4$-D35N*Mm*ACP spectrum (colored green) overlaid on the apo-D35N*Mm*ACP spectrum. B) $C_2$-D35NIacP spectrum (colored green) superimposed on the apo-D35NIacP spectrum (colored red). The spectrum was assigned based on previous assignments *(27)*. The figures were prepared using Sparky *(19)*.

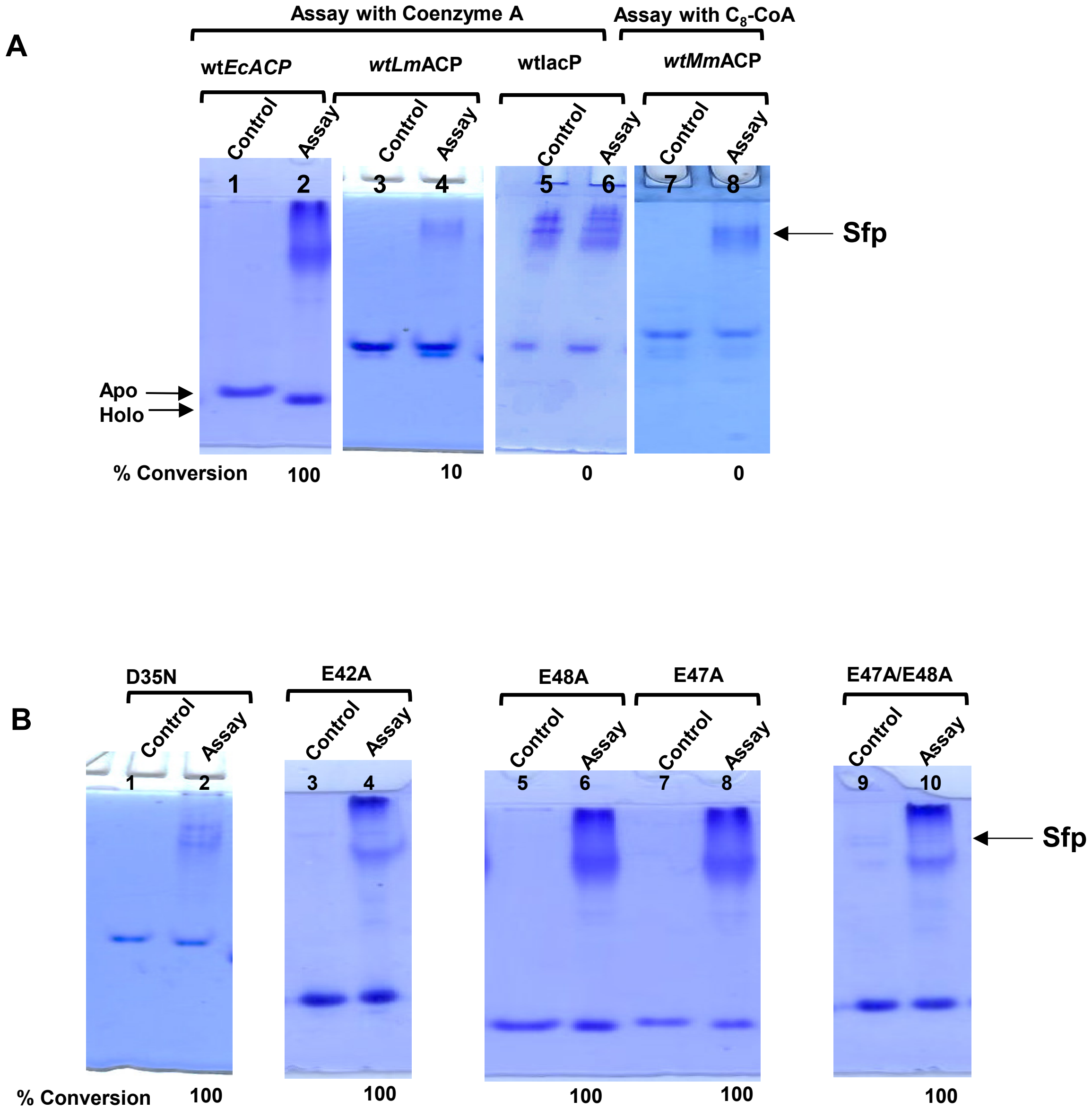
A
Assay with Coenzyme A
Assay with $C_8$-CoA
wt*EcACP*
*wtLm*ACP
wtlacP
*wtMm*ACP
Control
Assay
1
2
3
4
5
6
7
8
Sfp
Apo
Holo
% Conversion
100
10
0
0
B
D35N
E42A
E48A
E47A
E47A/E48A
9
10
100
100
100
100
100

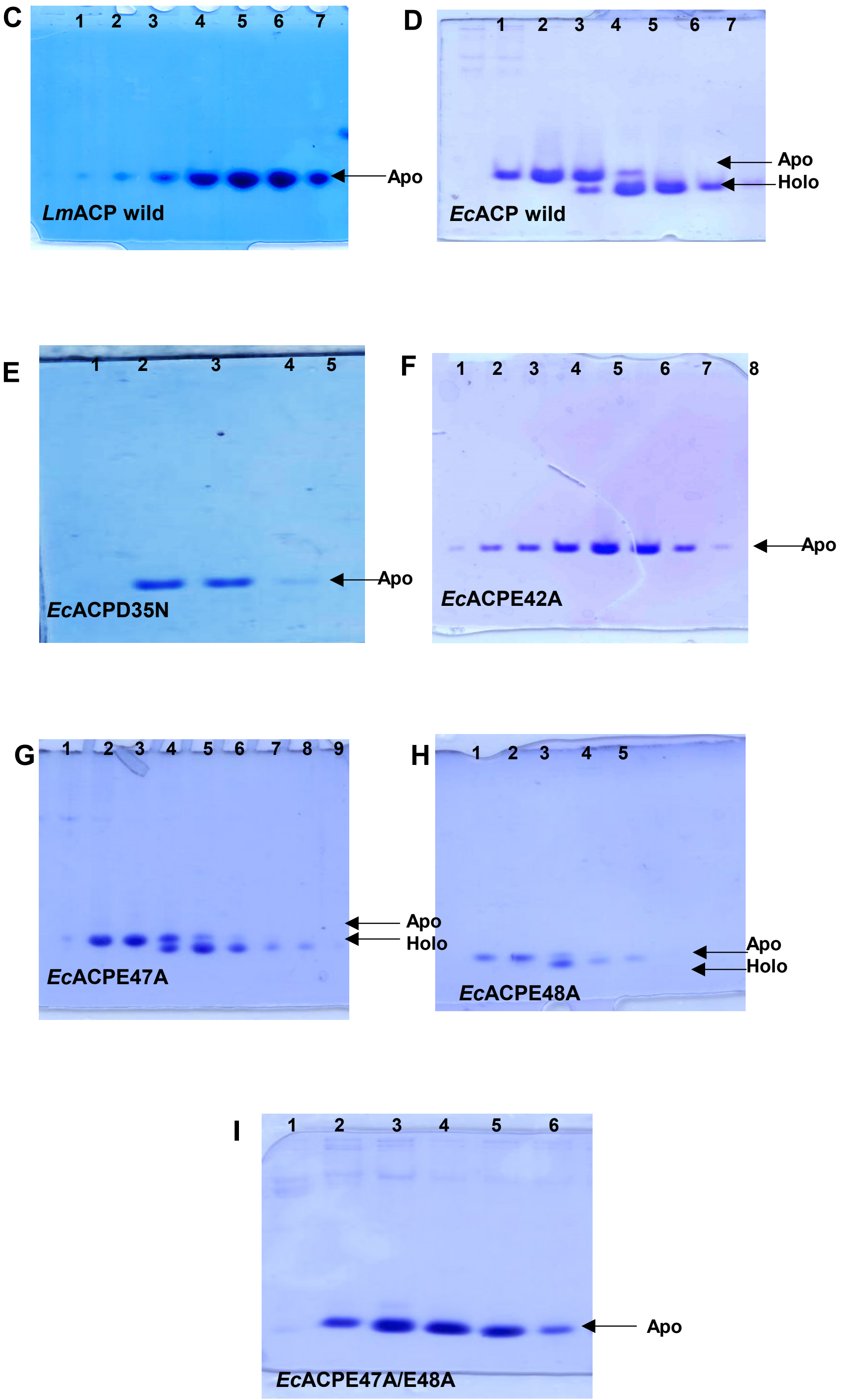
C
1 2 3 4 5 6 7
Apo
*Lm*ACP wild
D
1 2 3 4 5 6 7
Apo
Holo
*Ec*ACP wild
E
1 2 3 4 5
Apo
*Ec*ACPD35N
F
1 2 3 4 5 6 7 8
Apo
*Ec*ACPE42A
G
1 2 3 4 5 6 7 8 9
Apo
Holo
*Ec*ACPE47A
H
1 2 3 4 5
Apo
Holo
*Ec*ACPE48A
I
1 2 3 4 5 6
Apo
*Ec*ACPE47A/E48A

J

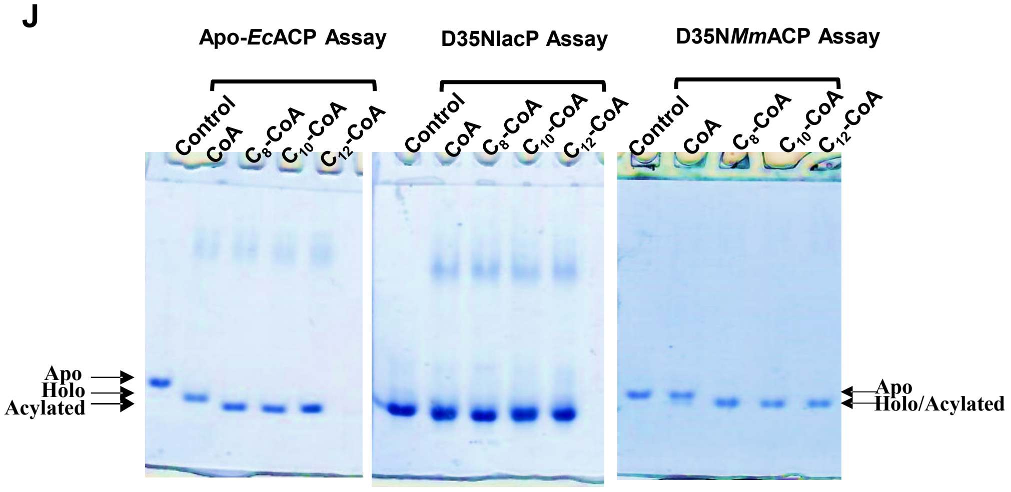


K

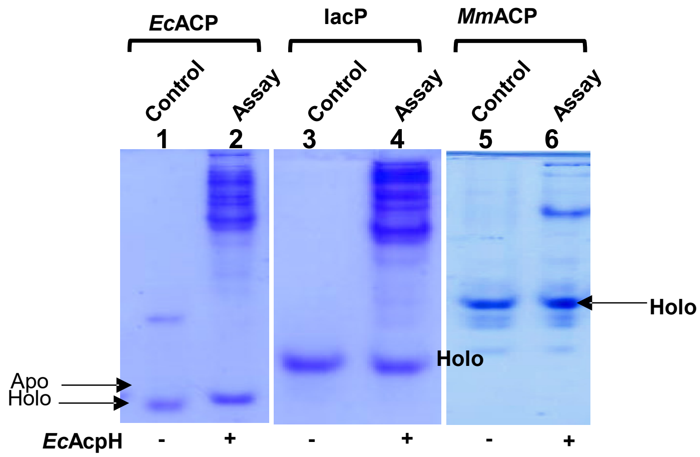


**Figure S4** 12% Native-PAGE gels reported in the main section. A & B) Full gels corresponding to Figures 2A &B. C-I) Gels reported in Figures 4A-G, H-I) Gels corresponding to Figures 6 A and B.
